# PAK1-dependant mechanotransduction enables myofibroblast nuclear adaptation and chromatin organisation during fibrosis

**DOI:** 10.1101/2023.03.31.535067

**Authors:** Elliot Jokl, Aoibheann F Mullan, Kara Simpson, Lindsay Birchall, Laurence Pearmain, Katherine Martin, James Pritchett, Rajesh Shah, Nigel W Hodson, Craig J Williams, Elizabeth Camacho, Leo Zeef, Ian Donaldson, Varinder S Athwal, Neil A Hanley, Karen Piper Hanley

**Affiliations:** Wellcome Centre for Cell-Matrix Research, Faculty of Biology, Medicine & Health, Manchester Academic Health Science Centre, University of Manchester, Oxford Road, Manchester, UK; Division of Diabetes, Endocrinology and Gastroenterology, Faculty of Biology, Medicine & Health, University of Manchester, Manchester Academic Health Science Centre, Oxford Road, Manchester, UK; Department of Life Sciences, Manchester Metropolitan University, Manchester, UK; Manchester University Hospital NHS Foundation Trust, Wythenshawe Hospital, Southmoor Road, Manchester, UK; BioAFM Facility, Faculty of Biology, Medicine & Health, University of Manchester, Manchester, UK; Department of Materials, University of Manchester, Manchester, UK; Division of Population Health, Health Services Research and Primary Care, School of Health Sciences, University of Manchester, Manchester, UK; Bioinformatics Core Facility Faculty of Biology, Medicine & Health, University of Manchester, Manchester, UK; Manchester University NHS Foundation Trust, Oxford Road, Manchester, UK

## Abstract

Myofibroblasts are responsible for scarring and organ stiffness during fibrosis. The scar propagates mechanical signals inducing a radical transformation in myofibroblast cell state linked to an increasingly pro-fibrotic phenotype. Here, we have discovered mechanical stress from progressive scarring induces nuclear softening and de-repression of heterochromatin. The parallel loss of H3K9Me3 enables a permissive state for distinct chromatin accessibility and profibrotic gene regulation. By integrating chromatin accessibility profiles (ATAC sequencing) we provide insight into the transcription network and open chromatin landscape underlying the switch in profibrotic myofibroblast states, emphasizing mechanoadaptive pathways linked to PAK1 as key drivers. Through genetic manipulation in liver and lung fibrosis, uncoupling PAK1-dependant signaling impaired the mechanoadaptive response in vitro and dramatically improved fibrosis in vivo. Moreover, we provide human validation for mechanisms underpinning PAK1 mediated mechanotransduction in liver and lung fibrosis. Collectively, these observations provide new insight into the nuclear mechanics driving the profibrotic chromatin landscape in fibrosis, highlighting actomyosin-dependent mechanisms linked to chromatin organisation as urgently needed therapeutic targets in fibrosis.

## Introduction

Fibrosis in any organ is characterised by the deposition of pathological extracellular matrix (ECM) leading to progressive scarring and ultimately causing loss of organ function. In the liver, fibrosis is potentially reversible during early stages. However, liver fibrosis is almost always diagnosed too late with limited treatment options. There is a massive unmet clinical need to halt liver fibrosis.

Activated hepatic stellate cells, (HSCs; liver specific myofibroblasts) are responsible for scar production during progressive liver fibrosis^1–5^. The physical scar increases tissue stiffness resulting in compromised tissue function that ultimately leads to liver failure^6–8^. Despite increasing knowledge of important signalling pathways promoting HSC activation and expression of key profibrotic components ^3, 5, 9–14^, many are also active in the healthy liver and by themselves, or even in aggregate, do not account for the complete phenotypic switch in cell state that distinguishes activated from quiescent HSCs. In recent years, we know that tissue stiffness can radically alter the cytoskeletal architecture of HSCs to influence cell behavior through cell surface integrin receptors ^3, 6, 8, 15^. For example, loss of integrin-β1 reduces the ability of HSCs to adhere firmly to pro-fibrotic extracellular matrix (ECM), with reduced actin organisation in the cytoskeleton ^3^. Downstream of integrin-β1, the serine/threonine-protein kinase P21-activated kinase group (PAK; associated with actomyosin signaling) and the transcriptional co-activator Yes-associated protein-1 (YAP1) are mechanosensing mechanisms responsible for myofibroblast activation and liver fibrosis ^3, 9^. Although informative, it is becoming increasingly evident that mechanical signals reach beyond the cytoskeleton toward the nucleus to induce a rapid pro-fibrotic physiological response ^16, 17^.

The nucleus is enclosed by the nuclear envelope (NE) composed of an outer and inner nuclear membrane ^17, 18^. The nuclear lamina (NL) is a lining on the inner nuclear membrane that contains lamin proteins, which provide elasticity to the NE allowing the nucleus to adapt to mechanical stimuli ^16–18^. The NE is connected to the cytoskeleton by the Linker of Nucleoskeleton and Cytoskeletal (LINC) complex of proteins, which also connects to the NL ^19^. Thus, the cytoskeleton, intimately linked to intracellular tension, is connected to the NL. Moreover, some genomic regions, called lamina-associated domains (LADs), are anchored to the NL ^20^. In mammalian cells, LADs can cover >30% of the genome and are generally associated with genome repression and heterochromatin marks such as lamina-associated Histone H3 tri-methyl Lysine 9 (H3K9Me3) ^21^. Thus, it becomes easy to conceive how cell tension, as increases during HSC activation, might alter spatial organisation of the genome and transcription ^18, 22^. This represents an under researched and emerging area for targeted anti-fibrotic therapy. However, to facilitate this research requires investigation in clinically relevant cellular models to avoid confounding factors associated with epigenetic mechanical memory from prolonged primary cell culture or use of cell lines.

Here, in freshly isolated liver HSCs, we begin from the perspective that liver myofibroblasts undergo a radical transformation in cell shape and state linked to an increasingly pro-fibrotic phenotype. For the first time our data identifies nuclear softening and de-repression of heterochromatin following H3K9Me3 loss as a permissive state for distinct chromatin accessibility and profibrotic gene regulation. We discovered this pro-fibrotic phenotypic switch could be reversed by uncoupling actomyosin signaling through *Pak1* deletion. In vivo, *Pak1*-loss during liver fibrosis improved scarring and blocked matrix-mediated myofibroblast activation. Through integrative analysis of RNA and Assay for Transposase-Accessible Chromatin (ATAC) sequencing data we provide detailed understanding of the transcriptional networks and open chromatin landscape underlying the switch in profibrotic myofibroblast cell states, emphasizing mechanoadaptive pathways linked to PAK1 as key drivers. As a core PAK1-dependant mechanism we provide further in vitro and in vivo evidence in models of lung fibrosis. Pathways capable of blocking or reversing this response represent highly desirable anti-fibrotic agents with implications across organ fibrosis.

## Results

### PAK1-dependant signaling promotes myofibroblast activation in human and mouse models of fibrosis

During liver fibrosis, increasing tissue stiffness induced by the pathological scar activates HSCs^3, 6, 8^. The in vitro model for activating HSCs on tissue culture plastic exemplifies this^3, 11, 14^. We have previously described PAK1 as the predominant group I PAK family member detected during HSC activation in vitro^3^. To understand the pathophysiological role of PAK1 in fibrosis, we initially modelled liver fibrosis in mice using carbon tetrachloride (CCl_4_) injections to induce parenchymal injury and bile duct ligation (BDL) for biliary injury. In both models, *Pak1* transcript was detected in elongated cells, characteristic of activated HSCs, localized to and within the fibrotic scar (Fig. 1a). *Pak1* transcripts were also evident in hepatocytes parallel to the scar (particularly during BDL induced fibrosis). These data are similar to our previous work indicating hepatocytes aligning the scar undergo a regenerative response indicative of the ductal plate during liver development ^9^. Similarly, in sections of human liver cirrhosis, PAK1 protein was robustly localized to PDGFRβ positive fibrotic areas indicating its relevance to human liver disease (Fig. 1b). Given these similarities across species, we next determine the importance of PAK1 in progression of liver fibrosis using our mouse model system to investigating the consequence of *Pak1*-loss during fibrosis. Liver weight and toxicity (assessed in serum by ALT and bilirubin levels) were unaltered following *Pak1*-loss (Supplementary Fig. 1). In livers from CCl4-induced fibrosis, fibrotic collagen deposition was significantly reduced in the hepatic parenchyma by picro sirius red (PSR) staining following *Pak1*-loss (Fig. 1c and d). Moreover, loss of *Pak1* also decreased the amount of alpha-Smooth muscle actin (α-SMA) staining in the liver implying reduced myofibroblast activation (Fig. 1c and e). Similarly, in BDL-induced liver fibrosis the marked peribiliary collagen deposition was greatly improved alongside reduced α-SMA staining and ductal hyperplasia (a feature of biliary injury shown by cytokeratin-19 (CK19) analysis) in *Pak1*-null animals (Fig. 2a-d). To determine the importance of PAK1 in broad mechanisms associated with myofibroblast activation in fibrosis, PAK1 was detected in regions of human lung fibrosis due to idiopathic pulmonary fibrosis (IPF; Supplementary Fig. 2a). As a second in vivo validation model and we carried bleomycin-induced lung fibrosis in *Pak1*-null animals. Similar to liver, compared to control mice, loss of *Pak1* significantly improved the extent of collagen deposition and fibrosis (Supplementary Fig. 2b-c). These data support PAK1 as a core regulator of myofibroblast activation during fibrosis and suggest mechanistic insight may highlight PAK1-dependant pathways as targets to improve fibrosis therapeutically.

**Figure 1.**
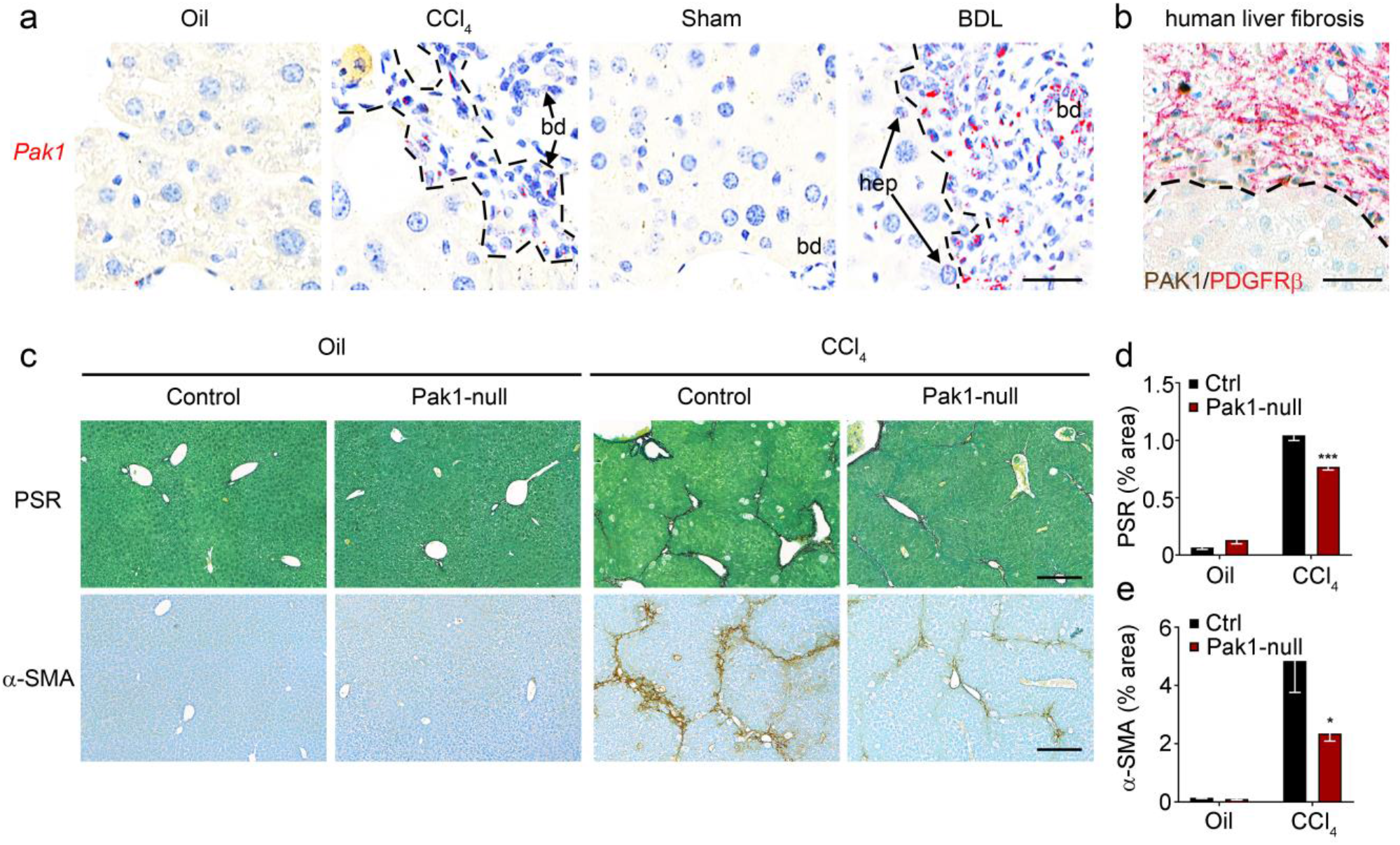
Pak1 is localized to fibrotic scar and its loss in vivo improves CCl_4_-induced liver fibrosis. (**a**) *Pak1* transcript (red) localized to cells within the fibrotic scar (hatched line) in both CCl_4_ and BDL-induced liver fibrosis in vivo. bd, bile duct indicated and hep, hepatocytes. Size bar = 25 µm. (**b**) PAK1 protein (brown) localized to the scar demarcated by PDGFRβ (red) and defined from hepatocytes via hatched line. Size bar = 25 µm. (**c**) PSR staining (collagen deposition in red) counterstained with fast green (top row) and α-SMA (brown) counterstained with toluidine blue (bottom) in olive oil (Oil) or chronic CCl4-induced fibrosis in control and Pak1-null mice (both n = 5). Size bar = 500 µm. (**d, e**) Quantification of surface area covered by PSR staining in **d** and α-SMA in **e** from representative image in **c**. Means ± SEM. *P<0.05, **P<0.005. Data analysed using 1-way ANOVA with Tukey’s post-hoc test (d, e).

**Figure 2.**
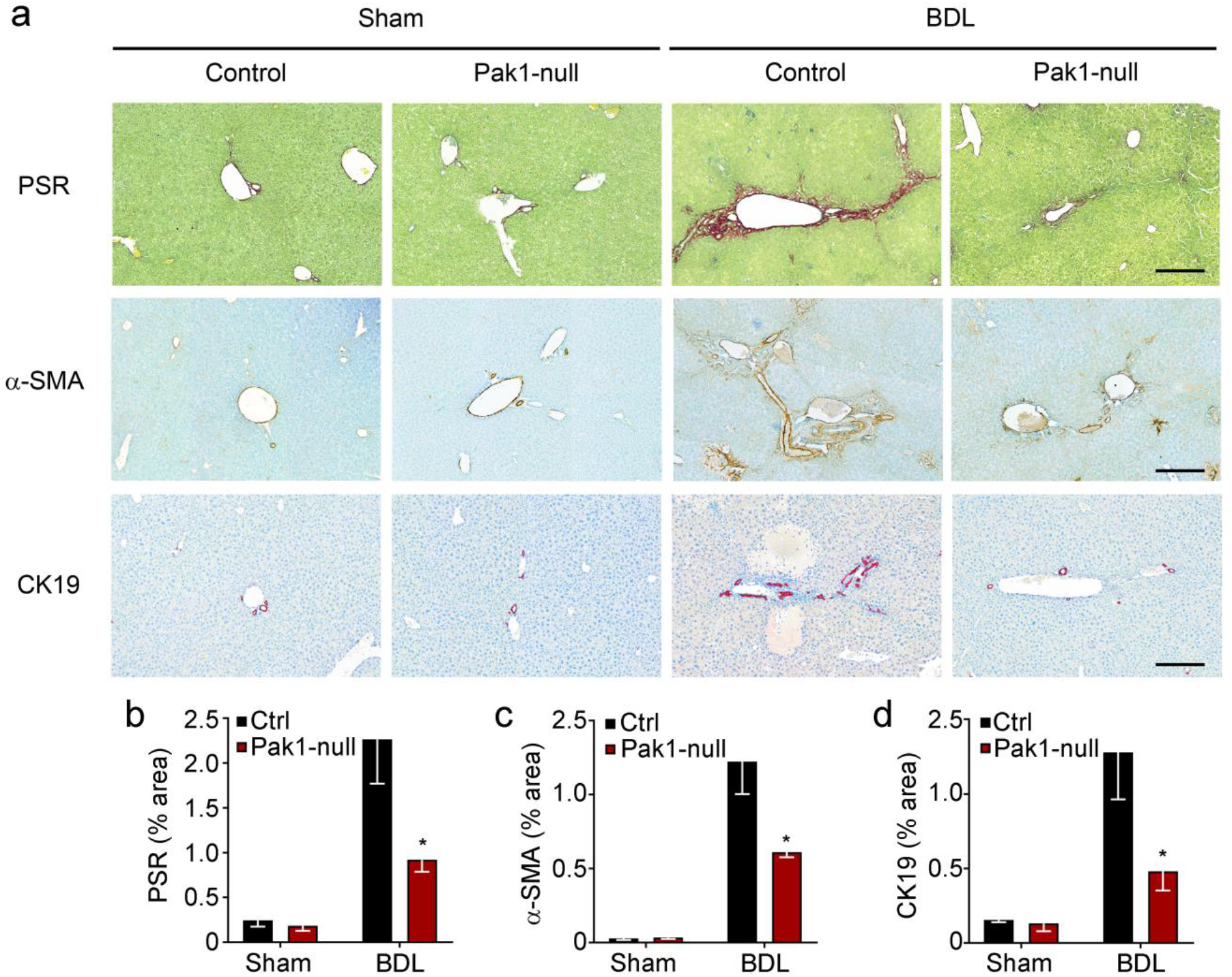
Pak1 loss improves biliary fibrosis in vivo. (**a**) PSR staining (collagen deposition in red) counterstained with fast green (top row), α-SMA (brown; middle row) and CK19 (brown; bottom row) counterstained with toluidine blue in sham operation (sham) or BDL-induced fibrosis in control and Pak1-null mice (both n = 5). Size bar = 500 µm. (**b-d**) Quantification of surface area covered by PSR staining in **b**, α-SMA in **c** and CK19 in **d** from representative image in **a**. Means ± SEM. *P<0.05. Data analysed using 1-way ANOVA with Tukey’s post-hoc test (b-d).

### Mechanical stress and profibrotic response is transmitted via PAK1 in liver myofiboblasts

To probe the mechanisms underlying PAK1 function in myofibroblasts, we extracted HSCs from wild type and *Pak1*-null livers and cultured on plastic for 7-10 days to model myofibroblast activation with the appearance of α-SMA, Sex determining region Y-box 9 (SOX9) and the major fibrotic ECM protein, Collagen type 1 (COL1) ^11, 13, 14^. Consistent with the in vivo data, following culture on plastic, *Pak1*-null activated HSCs had a reduction in the classical fibrotic markers α-SMA, COL1 and SOX9 (Supplementary Fig. 3a). Moreover, in line with our previous work^3^, *Pak1*-null HSCs displayed reduced amounts of integrin-β1, integrin alpha 11 (ITGA11) and focal adhesion kinase (FAK) proteins; suggesting impaired ECM sensing (Supplementary Fig. 3a). More broadly, transcriptomic analysis showed *Pak1* loss did not convert cells back to a quiescent phenotype but a largely intermediate state consistent with our previous work on HSCs lacking integrin-β1 (up-stream of PAK1; Supplementary Fig. 3b and Supplementary Data 1)^3^. To identify gene regulatory pathways responsible for the reduced profibrotic phenotype in *Pak1*-null myofibroblasts, we carried out hierarchical cluster analysis (Supplementary Fig. 3b) From eight clusters, we focused our attention to Cluster 5 with consistent reduction in differentially expressed genes following *Pak1*-loss in both quiescent and activated HSC populations (Supplementary Fig. 4). Significantly, the top twenty gene ontology terms were involved in contractility, ECM and cytoskeletal signaling (Supplementary Fig. 5).

In keeping with these data, stress fibers (marked by F-actin) were impaired and the myofibroblast-like appearance was lost in *Pak1*-null HSCs which had a rounded (more quiescent-like) phenotype (Fig. 3a). Similarly, *Pak1*-null cells displayed significantly reduced F-actin intensity compared to control cells using live cell imaging of fluorescently labelled endogenous F-actin (Fig. 3b and c); indicative of disrupted actin polymerization. *Pak1*-loss in activated myofibroblasts also disabled collagen gel contraction (Fig. 3d and e). To assess cell migration we used 24 hour single cell tracking of *Pak1*-deficient compared to control myofibroblasts. Although no marked difference in cell migration was detected in terms of total track length, the directionality of migration in *Pak1*-null HSCs was significantly perturbed, showing a more focal migratory pattern (Fig. 3f-i; Supplementary Figs. 6 and 7). To investigate this further, we explored whether the disrupted actin cytoskeleton impaired myofibroblast polarisation important for nuclear orientation in cell migration. Control and *Pak1*-null activated HSCs were plated onto fibronectin coated bow shaped micropatterns, to recapitulate patterning required in classical front-rear polarisation. Following culture and staining for the actin cytoskeleton, there was a noticeable failure in cytoskeletal organisation of cells lacking *Pak1* paralleled by perturbed nuclear orientation (Fig. 3j and k).

**Figure 3.**
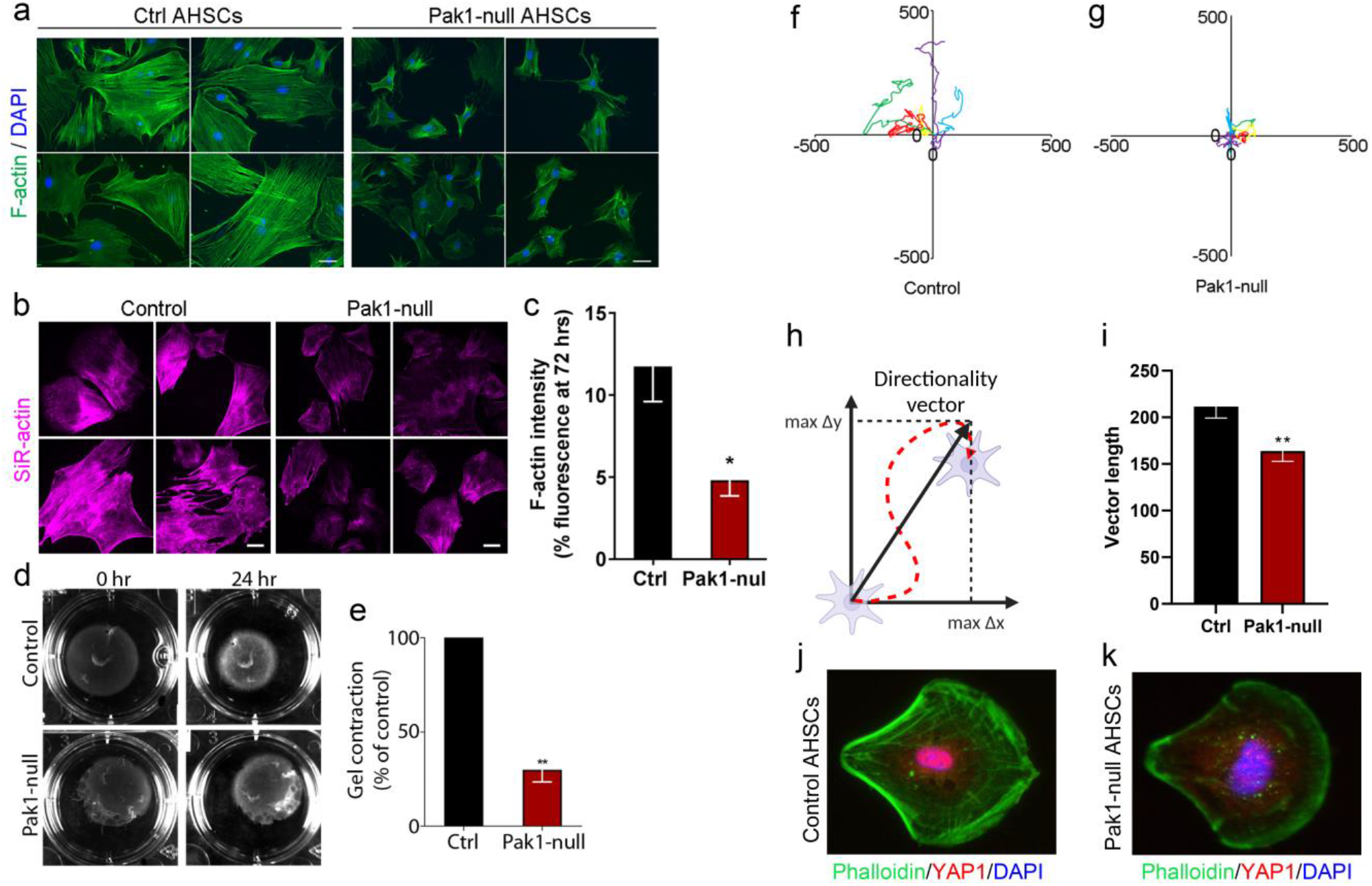
Functional characterization of liver myofibroblasts lacking Pak1. (**a**) Immunofluorescence staining for F-actin (green) and nuclei (DAPI; blue) in control and Pak1-null activated HSCs (AHSCs). (**b, c**) Live F-actin labelling in live control and Pak1-null AHSCs imaged at 72 hours. Example images are shown in **b** with fluorescence intensity quantification in **c** from three biological replicates with 15 cells per experiment analysed. (**d, e**) Contractile properties of control and Pak1-null AHSCs assessed by gel contraction assay. Representative image shown in **d** and quantification from three biological experiments shown in **e**. (**f-i**) Live cell migration over 24 hours from control and Pak1-null AHSCs. (**f, g**) Example track length shown (data from three biological replicates and 15 cell tracks). (**h, i**) Schematic demonstrating cell directionality assessed by the hypotenuse length of the triangle defined by the magnitude of movement in the x and y axes (**h**) and graphical representation (**i**). (**j, k**) Immunofluorescence for F-actin (green), YAP-1 (red) and nuclei (DAPI; blue) in control and Pak1-null AHSCs cultured on cross-bow micropatterns for 24 hours. Representative images of three independent experiments. Means ± SEM; n=3 biological replicates. *P<0.05, **P<0.01 by two-tailed t-test. Size bar = 5 µm.

### HSCs lacking Pak1 are unresponsive to their physical environment

Mechanical forces transmitted through the cell directly influence nuclear shape and function responsible for altered gene expression. To explore the mechanical behavior of activated HSCs we placed cells in a restricted physical environment and used YAP1 localisation as a mechanical readout. Following culture activation, control or *Pak1*-null HSCs were placed on fibronectin coated circular micropatterns. As expected, YAP1 localization in control HSCs was predominantly nuclear on large micropatterns (Fig. 4a). In contrast, *Pak1*-null HSCs displayed much reduced and limited YAP1 nuclear localization paralleled by a disrupted actin cytoskeleton marked by αSMA; indicative of impaired cell spreading and mechanical signaling (Fig. 4a).

**Figure 4.**
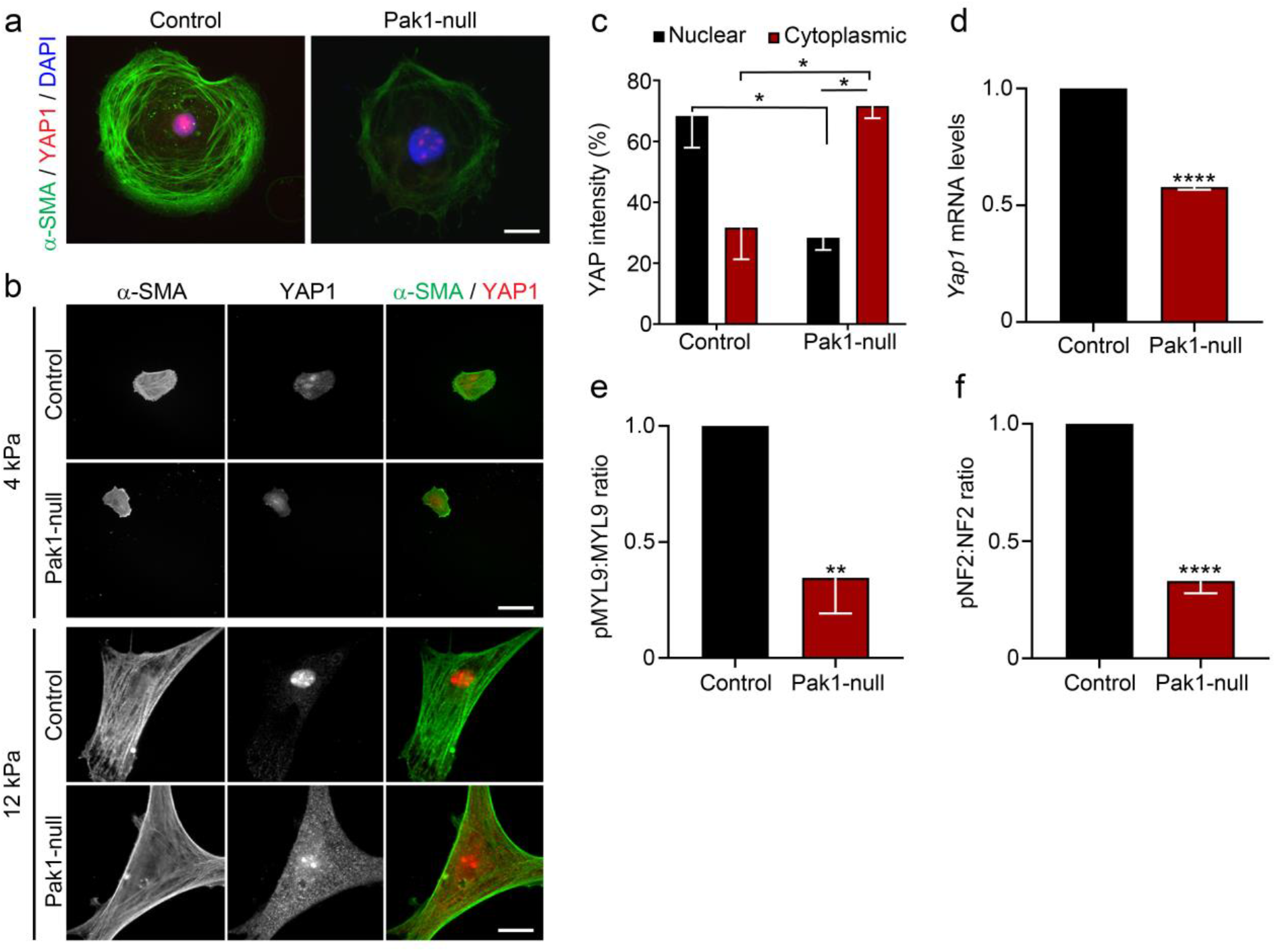
Mechanical response is impaired in Pak1-null liver myofiboblasts. (**a**) Immunofluorescence staining for α-SMA (green), YAP1 (red) and nuclei (DAPI; blue) in control and Pak1-null AHSCs cultured on circular micropatterns for 24 hours. Representative images of three independent experiments. (**b, c**) Immunofluorescence showing individual and merged images for α-SMA (green) and YAP1 (red) in control and Pak1-null AHSCs cultured on hydrogels at 4KPa and 12 KPa. Representative image in **b** and quantification in **c** showing the average intensity of nuclear and cytoplasmic YAP1 fluorescence of three biological replicate experiments from 15 cells analysed per experiment. (**d**) Quantification and relative levels of Yap1 mRNA in control and Pak1-null AHSCs. (**e, f**) Protein quantification and phosphorylation ratios shown for pMYL9 to total MYL9 (pMYL9:MYL9) in **e** and pNF2 to total NF2 (pNF2:NF2) in **f**. Means ± SEM; n=3 biological replicates. *P<0.05, **P<0.01, ****P<0.001. Data analysed using 1-way ANOVA with Tukey’s post-hoc test (c) and two tailed t-test (d-f). Size bar = 10 µm.

To provide insight from a physiologically relevant fibrotic environment, activated control or *Pak1*-null HSCs were cultured on hydrogels with differing stiffness to model physiologically normal liver elasticity (4kPa) or the fibrotic environment (12 kPa). Little difference in YAP1 localization was observed between control and *Pak1*-null HSCs at 4 kPa, however, interestingly, the profibrotic marker αSMA appeared more pronounced in control cells highlighting early cytoskeletal alterations following *Pak1*-loss (Fig. 4b). HSCs cultured on 12 kPa maintained the altered αSMA appearance, whereas YAP1 nuclear localization was greatly diminished in Pak1-null cells (Figs. 4b and c). In support, Pak1-null cells displayed reduced RNA levels of *Yap1* and diminished phosphorylation of the direct downstream target MYL9 (Figs. 4d and e); important in promoting actomyosin interactions and force generation. Mechanistically, PAK1 has been linked to Merlin/NF2 phosphorylation and subsequently alleviating its negative regulation of YAP1^23^. In line with this, activated HSCs lacking *Pak1* have reduced NF2 phosphorylation (Fig. 4f).

### Myofibroblast activation induces a radically altered cell state associated with nuclear adaptation

Collectively our data indicated myofibroblasts lacking PAK1 were less fibrotic and unresponsive to their physical environment through perturbed cytoskeletal and mechanosignalling. To explore how the physical environment alters myofibroblast cell state, we modelled the increasing pro-fibrotic response in culture activated HSCs at 1, 3, 5, 7 and 10 days. In addition to the characteristic cytoskeletal shape change (Supplementary Fig. 8a)^3, 11, 14^, we uncovered nuclear size was increased as HSCs activate (Fig. 1 a-b; Supplementary Fig. 8a-b). In parallel, DAPI intensity was noticeably reduced suggesting altered chromatin organisation (Fig. 1c; Supplementary Fig. 8a-b). Following exponential regression analysis, there was a statistically significant relationship suggesting nuclear area explains 96 % of the variance in DAPI intensity (Fig. 1c; R^2^ = 0.96; p <0.001). These data were similarly recapitulated in culture activated lung fibroblasts, suggestive of similar state changes at the cellular level in lung, potentially indicating broad function in organ fibrosis (Supplementary Fig. 9). To explore the impact of this mechanoadaptive, response we used atomic force microscopy (AFM) to assess the physical properties of the nucleus and discovered activated myofibroblasts have reduced nuclear stiffness in response to a stiff environment (Fig. 1d).

To understand this adaptive response further, we used our physiologically relevant *Pak1*-null HSC model to uncouple actomyosin and, in part, alleviate the mechanical stress response. We first focused on the composition of the nuclear envelope. Lamina proteins, including Lamin A/C (LMNA/C) and Lamin B1 (LMNB1), line the inside of the nuclear envelope where disruption of their function and composition can lead to altered nuclear morphology and gene expression^24^. Increased LMNA/C has previously been associated with cells cultured on stiff matrices^25^. Whereas, from our genomic datasets, *LmnA* and *Lamb1* are both highly expressed in activated HSCs (Supplementary Fig. 10a-d; Supplementary data 1). In keeping with this, although very little difference in LMNA/C protein levels were detected (Fig. 5e), LMNB1 was almost double in *Pak1*-null HSCs (Fig. 5f). Using AFM we tested how these changes related to nuclear stiffness. Compared to control HSCs, loss of *Pak1* induced nuclear stiffening (Fig. 5g-h). In parallel, culturing activated *Pak1*-null HSCs on hydrogels decreased nuclear area at physiologically relevant liver stiffness; consistent with an impaired mechano-response (Figs. 5i-j). These data were reminiscent of a quiescent phenotype observed in our profile of HSC activation over time (Fig. 5a) further emphasizing the critical role of PAK1-dependant signaling in the profibrotic response of myofibroblasts.

**Figure 5.**
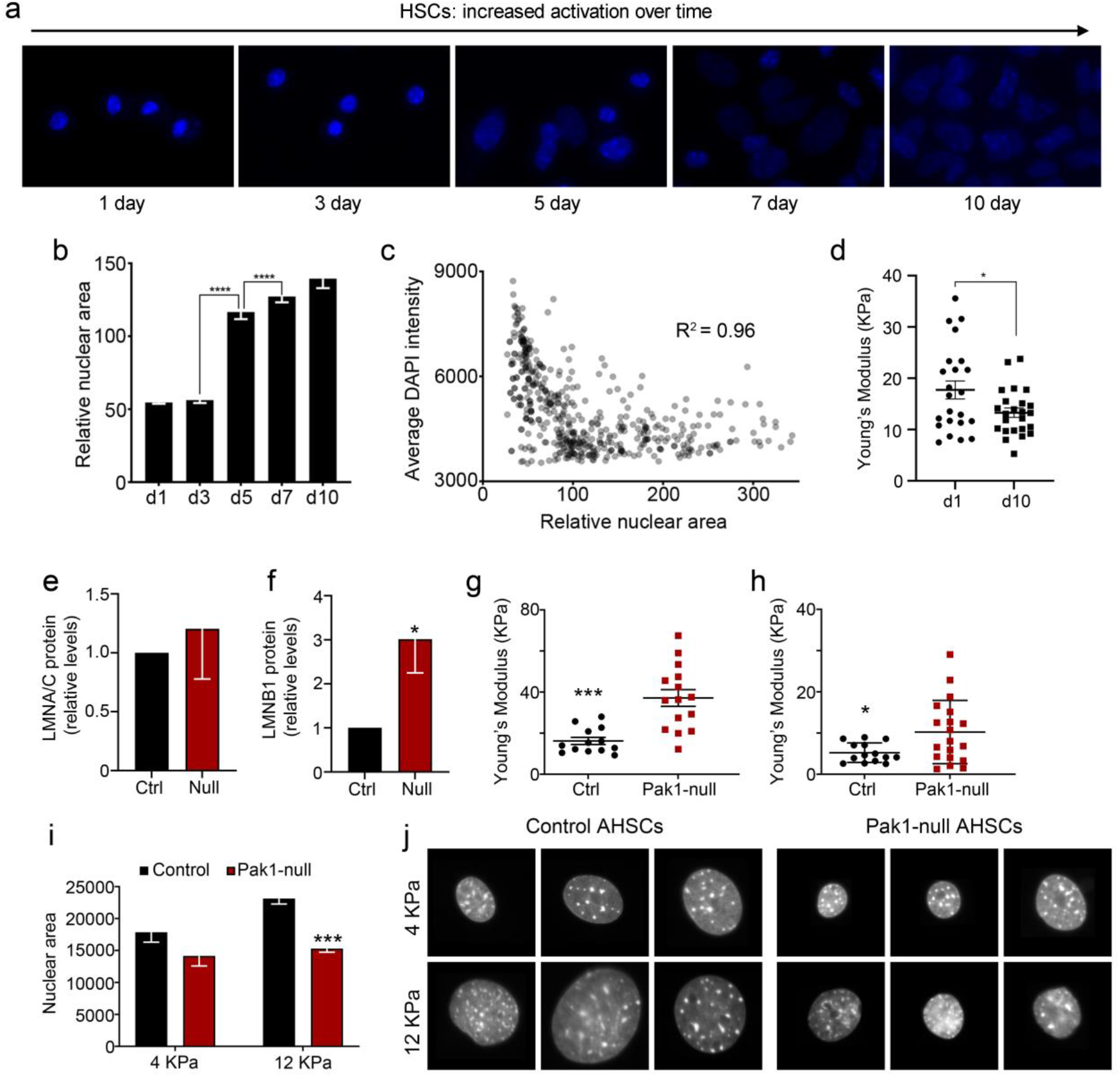
Altered cell state during liver myofibroblast activation. (**a-b**) Representative DAPI (blue) images of HSC nuclei in **a** and quantification of nuclear area during HSC activation on plastic over time in days from quiescent (d1) to activated (d10) in **b**. (**c**) Graphical representation of all cells from **b** and inverse correlation of average DAPI intensity with nuclear size (R^2^ = 0.96; P <0.001).). (**d**) Nuclear stiffness measurements displayed as KPa using AFM in quiescent (d1) and activated (d10) HSCs. (**e, f**) Protein quantification of LMNA/C in **e** and LMNB1 in **f** in control and Pak1-null AHSCs. (**g, h**) Nuclear stiffness measurements displayed as KPa using AFM in fixed in **g** and live in **h** control and Pak1-null AHSCs. (**i, j**) Quantification in **i** and representative DAPI images in **j** of nuclear area in control and Pak1-null AHSCs cultured on hydrogels at 4KPa and 12 KPa. Means ± SEM; n=3 biological replicates. *P<0.05, ***P<0.005 and ****P<0.001. Data analysed using 1-way ANOVA with Tukey’s post-hoc test (b) and two tailed t-test (d-i).

### H3K9Me3-loss promotes nuclear remodelling and chromatin organisation in myofibroblasts during fibrosis

We next wanted to understand how cell state changes at the nucleus impacted on chromatin organization and the phenotypic profibrotic gene expression. Mechanistically, we observed loss of the heterochromatin marker, H3K9Me3 as HSCs activate (Fig. 6a). Similarly, protein expression of H3K9Me3 associated methyltransferase, Suppressor of variegation 3-9 Homolog 1 (SUV39H1), was also decreased (Figs. 6b). In mouse models of liver fibrosis, H3K9Me3 maintained robust expression in hepatocytes and bile ducts, whereas expression in myofibroblasts displayed lower H3K9Me3 intensity within the vimentin (VIM) positive nuclei associated with the scar compared to the surrounding tissue (Fig. 6c-d; Supplementary Fig. 11a). Significantly, this altered nuclear cell state parallels a pro-fibrotic switch in gene expression. From single cell transcriptomics studies^26, 27^, core signatures associated with quiescent and activated HSCs were faithfully recapitulated in our genomic datasets (Supplementary Fig. 10a-d; Supplementary Data 1). Whereas, signatures associated with collagen and ECM modifications were highly induced alongside mechanotransduction pathways (Figs. 10 c-d), emphasizing the environment and intracellular response as critical mediators of the observed physiological response at the nucleus. To explore this further and understand how mechanical stress promotes H3K9Me3 heterochromatin function and nuclear properties of myofibroblasts, we analyzed the levels of H3K9Me3 and the methyltransferases SUV39H1 following *Pak1* loss. Following HSC activation we uncovered increased levels of H3K9Me3 and SUV39H1 in *Pak1*-null compared to control HSCs indicative of a repressed chromatin state, reminiscent of quiescent HSCs (Fig. 6e-g). To further validate the link between H3K9Me3 methyltransferase activity and HSC activation, simultaneous siRNA knockdown of *SUV39H1* and *SETDB1* in the LX2 HSC human cell line resulted in elevated *COL1A1* and *ACTA2* transcript, consistent with a more activated state (Fig. 6h).

**Figure 6.**
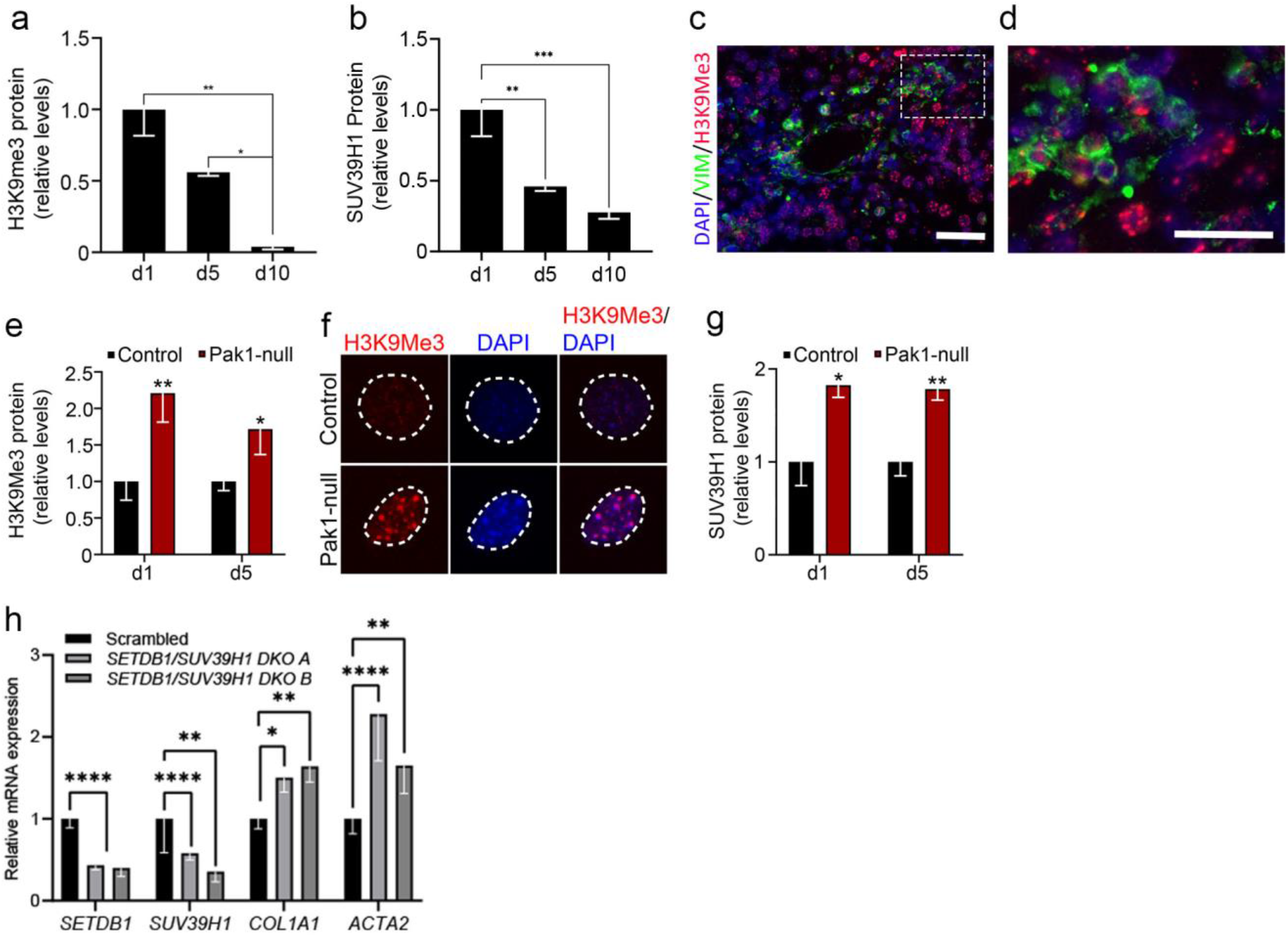
H3K9Me3 loss is associated with profibrotic myofibroblasts. (**a, b**) Quantification of H3K9me3 and SUV39H1 proteins during HSC activation. (**c, d**) Immunofluorescence showing lower level H3K9Me3 (red) associated with VIM (green) positive myofibroblasts in sections of murine liver fibrosis; DAPI (blue). Boxed area in **c** magnified in **d**. (**e, f**) Protein quantification of H3K9Me3 in **e** and representative immunofluorescence staining for H3K9Me3 (red) and nuclei (DAPI; blue) in **f** in control and Pak1-null AHSCs. Hatched white line in **f** indicates external surface of nucleus. (**g**) Quantification of SUV39H1 protein in control and Pak1-null AHSCs. (**h**) Increased RNA levels of profibrotic markers *COL1A1* and *ACTA2* following abrogation of *SETDB11* and *SUV39H1* methyltransferases. Two independent siRNA are shown (A and B) relative to their respective scrambled control levels. Means ± SEM; n=3 biological replicates. *P<0.05, **P<0.01, ****P<0.001. Data analysed using 1-way ANOVA with Tukey’s post-hoc test (a, b, h) and two tailed t-test (e, g). Size bar = 20 µm.

### Gene de-repression following H3K9Me3 loss correlates with a distinct profibrotic gene-regulatory network

To investigate the chromatin accessibility and distinct gene regulatory networks associated with profibrotic myofibroblasts, we analysed the genome-wide distribution of H3K9Me3 in the two distinct states of liver HSCs, quiescent and activated using CUT&TAG. Following sequencing we characterized the dynamic distribution of H3K9Me3 peaks from two distinct antibodies. Consistent with our in vitro and in vivo data, using MACS2 peak calling, genome wide enrichment of H3K9Me3 was detected in qHSCs compared to aHSCs (Supplementary Fig. 11b-c). As further validation, we used Sparse Enrichment Analysis (SEACR), a peak caller method designed for processing CUT&TAG data. Similar to MACS2, SEACR identified an average of 15,666 total peaks in quiescent compared to only 3,113 in activated HSCs across the two independent antibodies. We assumed this would correlate with repression of activation associated genes, however, the H3K9Me3 enrichment observed in qHSCs appeared indiscriminately distributed between promotors of both quiescence and activation associated genes (Supplementary Fig. 11d). These data suggested that although H3K9Me3 marks alone may not provide bespoke gene silencing in HSCs, de-repression of heterochromatin following H3K9Me3 loss provides a permissive state for distinct chromatin accessibility and profibrotic gene regulation in aHSCs. To further understand how chromatin reorganization correlates with gene-regulatory networks, we carried out parallel ATAC–seq and RNA–seq on quiescent and activated HSCs. Candidate regions of open chromatin ‘peaks’ were identified using MACS2. Using differential binding analysis, chromatin accessibility profiles showed cell state specificity with gene expression. Significantly, aHSCs displayed a greater number of ATAC-seq candidate peaks, implying chromatin in aHSCs was more globally accessible (Fig. 7a), consistent with H3K9Me3 loss.

**Figure 7.**
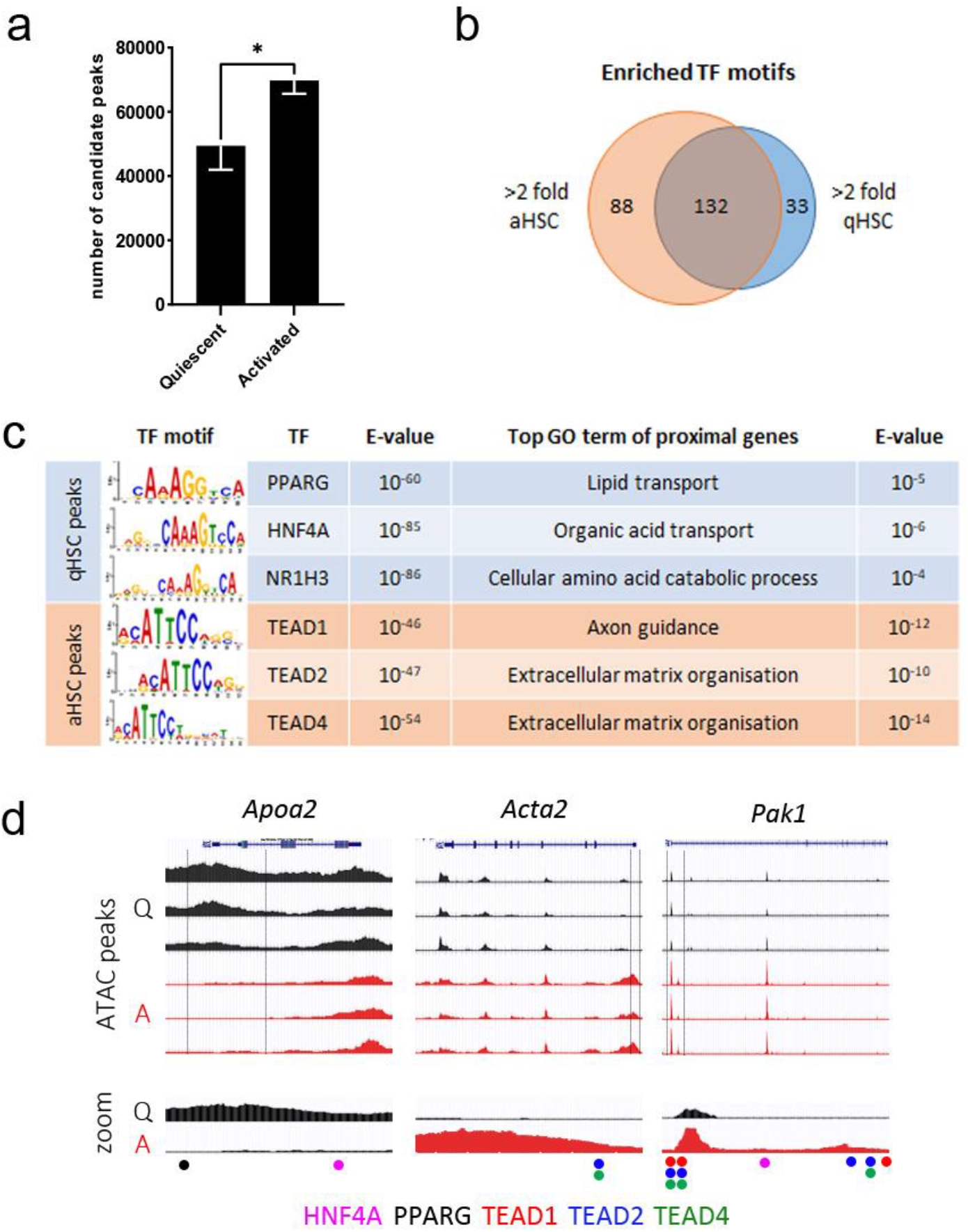
Chromatin organization correlates with profibrotic gene-regulatory networks. (**a**) Significantly higher number of candidate ATAC peaks in activated cells. (**b**) Venn diagram of TF motifs identified by Simple Enrichment Analysis of ATAC peaks with >2-fold higher concentration in aHSCs or qHSCs. 88 motifs unique to aHSCs, 33 unique to qHSCs and 132 common to both. (**c**) Selected TF motifs along with e-value for enrichment and the top GO Biological Function returned based on the list of proximal genes to motif-containing peaks. (**d**) ATAC tracks showing chromatin accessibility in quiescent (Q) and activated (A) HSCs. *Apoa2* shows increased accessibility in the quiescent state, conversely *Acta2* and *Pak1* show increased accessibility in the activated state. Indicated motifs from selected TFs from **c** in zoom image (bottom tracks). Means ± SEM; n=3 biological replicates. *P<0.05, by two-tailed t-test.

To determine the transcriptional networks underlying the quiescent and activated HSC states we initially undertook Simple Enrichment Analysis of TF motifs in the sequences of all ATAC peaks which showed >2 or <-2 -fold peak concentration in the activated HSCs compared to quiescent. This was followed by gene ontology analysis of genes proximal to ATAC peaks. As expected, many motifs were common to both quiescent and activated HSC states, however, 88 were unique to the activated state, including TEAD1, TEAD2 and TEAD4 and 33 were unique to the quiescent state, including PPARG, NR1H3 and HNF4A (Fig. 7b-c; Supplementary data 1). Indeed, HNF4A and PPARG sites are evident around the promotor of *Apoa2*, which is more open in the quiescent state (Fig. 7d). Similarly, TEAD2 and TEAD4 sites are evident around the promotor of *Acta2* which is more open in activation (Fig. 7d). Significantly and in keeping with our data suggesting PAK1 is critical mediator of mechanoadaptive pathways in HSC states, several TEAD sites were present around the promotor of *Pak1* (Fig. 7d).

### Understanding the transcription factor networks driving myofibroblast cell state

Further integration of our RNA and ATAC-seq data provided a more detailed understanding of how gene expression is regulated in each HSC state (Supplementary Fig. 12a). Focusing on ATAC peaks at gene promotors, we used sequential filtering to generate gene lists based on significant alteration of promotor peak concentration and RNA expression in each state (Supplementary Fig. 12b; Supplementary data 2). Gene Ontology enrichment identified genes with a more open chromatin profile and upregulated in activated HSCs (OAUA) were enriched for ECM organization, in line with their pro-fibrotic role. Whereas, genes more open and upregulated in quiescent HSCs (OQUQ) displayed enrichment for lipid metabolism, recapitulating classical quiescent phenotype (Supplementary Fig. 12c). Interestingly, the transcription factors *Gfi1*, *Sall1* and *Zeb1* associated with establishing H3K9me3-based heterochromatin domains were increased in the quiescent state compared to activated HSCs (Supplementary Table 1). Moreover, *Sall1* appeared more accessible with increased regions of open chromatin in qHSCs (OQUQ; Supplementary Table 1).

By combining motif enrichment at promotor peaks with respective RNA expression, we generated a regulatory TF network associated with the quiescent and activated-HSC state. This highlighted HFF4A, IRF7, STAT1/2 among others (Supplementary Fig. 13a) as enriched in the promotors of OQUQ genes and expressed more highly in the quiescent state. Similarly, ATF3, FOSL2 and CREB3/5 were amongst those enriched in the promotors of OAUA genes and expressed more in the activated state. (Supplementary Fig. 13a). Interestingly, overlaying our PAK1-null qHSC genomic data set with these defined TF networks showed significantly higher expression of several quiescent associated TFs, including STAT1/2 and FLI1 (Supplementary Fig. 13b-c; Supplementary Tables 1-2). These results provide further evidence that PAK1-null HSCs have a more quiescent transcriptional state, which may underlie their reduced mechanosensitivity and fibrotic potential observed in our in vitro and in vivo experiments. FLI1 in particular has been shown to suppress *Col1* expression, and in the context of ewing sarcoma (a highly metastatic bone cancer) disrupts YAP1/TEAD mechanotransduction^28, 29^.

Collectively, this study provides previously unappreciated insight into nuclear mechanics driving pro-fibrotic liver myofibroblasts and highlights PAK1-dependent regulatory mechanisms linked to chromatin state as novel targetable areas for urgently needed therapeutic targets in fibrosis.

## Discussion

The cellular process underlying fibrosis in any organ is broadly similar^30^. In the liver, regardless of the source, failure to resolve liver injury leads to progressive fibrosis^31^. Activated HSCs are widely regarded as the liver myofibroblasts responsible for tissue damaging ECM deposition in fibrosis where the mechanical environment, in part, perpetuates this response^3, 5, 8, 9^. As a general principle, nuclear stiffness scales with tissue stiffness where brain cells, for example, would have a softer nucleus that bone^25^. However, the response of cells within an organ and at the multicellular level is likely to be more complex, particularly in disease states.

Tissue stiffness is a feature of fibrotic organs^8, 9^. Extent of scarring and stiffness in liver is a diagnostic determinant of disease severity^8, 9, 32^. In light of this and previous studies, we might predict quiescent HSCs, as exist in normal healthy liver to have a softer nucleus than activated HSCs, as exist in fibrotic liver. However, here we discover that when HSCs encounter a stiffer environment the nuclear response is to soften. Significantly, this is in parallel to a radical shift in cytoskeletal cell shape and pro-fibrotic gene expression.

The reasons underlying this are likely two fold. First, pro-fibrotic HSCs are contractile, migratory myofibroblasts and as part of this phenotype a malleable nucleus would promote their ability to infiltrate the surrounding tissue and further deposit ECM. Second, although activated HSCs have been associated with promoting an environment conducive to cancer in end stage cirrhosis^33^, HSCs themselves are not known to be associated with DNA damage. In keeping with studies in skin^22^, nuclear softening would provide a protective mechanism and propagate the pro-fibrotic response.

It is clear that the shift in cell shape from quiescent to activated HSCs, and increasingly profibrotic myofibroblasts in lung, is paralleled by a major change in the actin cytoskeleton^3, 9–11, 13, 14^. Through uncoupling the actin cytoskeleton by *Pak1*-loss we highlight a functional mechanoresponse of liver myofibroblasts driven by nuclear deformation and H3K9me3-mediated chromatin remodeling. These findings are consistent with a profound switch in profibrotic gene expression and one that further permits their migratory, contractile phenotype.

Interestingly, during liver development loss of H3K9me3 correlates with hepatocyte differentiation and maturation^34^. More broadly, although the role of H3K9me3 in gene silencing is complex, a number of studies have indicated its increasingly important function in cell-type specific chromatin organization and gene expression^21, 34–36^. H3K9me3 is associated with condensed chromatin and restricted TF binding due to reduced accessibility of heterochromatic promoters^36–39^. Similarly, in this study, using ATAC-seq, we demonstrated increased chromatin accessibility parallels loss of H3K9me3. The correlation with enhanced nuclear adaptation, measured by AFM, and pro-fibrotic gene expression as quiescent HSCs switch to an activated phenotype supports the idea that H3K9me3 compacts chromatin structure and restricts gene expression programmes. These data have similarities to other studies and supports the idea that changes in H3K9me3 deposition provides a distinct chromatin structural organization permissible to tissue specific TF networks^21, 34, 36–39^.

Linked to these findings, we also observed altered expression of lamin proteins, particularly Lamin B, following impaired actin signaling. Aside from their structural properties as components of the inner NE, lamins are also associated with chromatin organization and gene expression though LADs^20^. Whether the relative composition of lamin proteins in HSCs also plays a role in creating permissible gene regulatory environments through their different expression and nucleoplasmic localization is unknown. Uncovering this may provide insight into the 3D chromatin structure and fine control mechanisms of HSCs and transformation into pro-fibrotic myofibroblasts. For cell specific therapeutic targeting, a comprehensive understanding of external mechanical cues through to signals that impact nuclear function seem critical to identifying pharmaceutical targets to block or reverse fibrosis^40^. As a first step to translation, we provide evidence that targeting actomyosin signaling during liver and lung fibrosis in vivo significantly reduces scarring and liver myofibroblast accumulation. However, our collective data also highlight H3K9Me3 as a potential therapeutic antifibrotic target in liver disease through modulation of methyltransferases (e.g. SUV39H1)^40^. In other models of organ fibrosis, although not assessed in a cell specific manner, increase levels of H3K9me3 have been highlighted in kidney fibrosis^41^. More interestingly, this study did show a more diffuse spatial distribution of H3K9me3 in response to TGF-β (the main profibrotic cytokine in organ fibrosis) which parallels our observation of chromatin decondensing during in vitro activation. Overall, given mechanisms underlying fibrosis are common across organs, our study and datasets have broad appeal across chronic disease where fibrosis is a progressive step to organ dysfunction.

## Methods

### Animals

The *Pak1*-null mice were developed by Dr Chernoff (Fox Chase Centre, Philadelphia) and gifted by Dr Solaro (University of Illinois) and Dr Chernoff^42^. Animals were genotyped by PCR with a 280bp wild-type product detected with forward 5’-GCCCTTCACAGGAGCTTAATGA-3’ and reverse 5’-GAAAGGACTGAATCTAATAGCA-3’ and a 360bp Pak1-null product detected using the same forward primer and reverse 5’-CATTTGTCACGTCCTGCACGA-3’. Animals were housed, maintained, and experiments performed in accordance to UK Government Home Office regulations and under approval from the University of Manchester Ethical Review Committee.

### Primary cell isolation and culture

Mouse hepatic stellate cells (HSCs) were isolated as previously described^3, 9, 11^. Briefly, livers were perfused with HBSS containing collagenase and pronase via the portal vein prior to mechanical disruption of tissue and isolation of HSCs via an Optiprep density gradient (Sigma). Cells were cultured in high glucose DMEM enriched with L-glutamine, pen-strep and 16% serum. Mouse lung fibroblasts were isolated by incubating lung tissue minced using razor blades in a collagenase/pronase solution at 37 degrees for approximately 1 hour with periodic agitation, followed by straining through a 40 micron filter, pelleting at 4 degrees at 800g for 5 minutes and resuspension in high glucose DMEM enriched with L-glutamine, pen-strep and 10% serum before plating.

### Animal models of fibrosis

Bile Duct Ligation (BDL) and carbon tetrachloride (CCl_4_) models of fibrosis were performed as previously described on *Pak1*-null and WT (FVB) control animals^3, 9^. CCl_4_ was administered twice weekly by intraperitoneal (i.p.) injections of 2μl per g body weight CCl_4_ (Sigma) at ratio of 1:3 by volume in olive oil (Sigma) or olive oil alone (control) twice weekly for 8 weeks. BDL was carried out under anaesthesia using two ligatures to ligate the bile duct. Sham-operated mice underwent the same procedure but without tying the ligatures. BDL mice developed cholestasis and associated fibrosis over a 14-day period. For bleomycin induced lung fibrosis, 2.5IU per kg body weight of bleomycin in saline, or an equivalent volume of saline alone, was delivered by intratracheal administration under anaesthesia. These mice developed lung fibrosis over a 21 day period.

At the end of these experiments, animals were sacrificed, blood taken for liver function biochemistry, liver and body weight recorded and tissues processed for histology.

### Clinical samples

Liver tissues were obtained with informed consent and ethical approval (National Research Ethics Service, REC 14/NW1260/22) from the Manchester Foundation Trust Biobank. Human lung sections were obtained following informed consent and ethical approval (National Research Ethics Service, REC 20/NW/0302) from the Manchester Allergy, Respiratory and Thoracic Surgery (ManARTS) Biobank (study number M2014-18).

### siRNA knockdown

*SUV39H1* and *SETDB1* genes were targeted with FlexiTube GeneSolution siRNA (Qiagen). Two independent siRNA pools were generated, each containing a pair of siRNAs against each gene: pool 1 (Hs_SETDB1_2, Hs_SETDB1_7, Hs_SUV39H1_1, Hs_SUV39H1_2) and pool 2 (Hs_SETDB1_8, Hs_SETDB1_9, Hs_SUV39H1_4, Hs_SUV39H1_7). siRNAs, or an equivalent amount of AllStars negative control siRNA (Qiagen) were transfected into LX2 cells using Lipofectamine 3000 transfection reagent as per manufacturer’s instructions. Cells were harvested at 48hrs post transfection prior to RNA extraction and qPCR analysis as described below.

### Immunoblotting and quantitative RT-PCR

Protein from cells and tissues was extracted in RIPA buffer and expression levels determined by a standard western blotting protocol using antibodies with total protein on the transfer blot or B-actin used as a loading control as indicated in the main text. The following primary and secondary antibodies were used: anti-ITGB1 (MAB1997, Millipore, dilution: 1:1000), anti-α-SMA (NCL-L-SMA, Leica Biosystems, dilution: 1:500), anti-COL1 (1310-01, Southern Biotech, dilution:1:1000), anti-SOX9 (AB5535, Millipore, dilution: 1:5000), anti-MYL9 (3672, Cell Signalling, dilution: 1:500), anti-YAP (Sc-271134, Santa Cruz, dilution: 1:1000), anti-PMYL9 (#3671P, Cell Signalling, 1:500), anti-PAK1 (#2602, Cell Signalling, dilution: 1:1000), anti-ITGA11 (MAB4235, R & D Systems, dilution: 1:500), anti-FAK (CST/3285, Cell Signalling, 1:1000), anti-PNF2 (CST/9163, Cell Signalling, 1:500), anti-NF2 (HPA003097, Atlas, 1:500), anti-LMNA/C (CST/2032, Cell Signalling, 1:1000), anti-LMNB1 (374015, Santa Cruz, 1:1000), anti-H3K9Me3 (CST/13969, 1:1000), anti-SUV39H1 (CST/8729, Cell Signalling, 1:500), anti-SETDB1 (ab12317, Abcam, 1:1000), anti-β-actin HRP conjugate (A3854, Sigma, dilution: 1:50000) and species-specific HRP conjugated secondary antibodies (GE Healthcare).

RNA was extracted using a RNeasy purification kit (Qiagen) followed by cDNA conversion using High Capacity RNA-to-cDNA kit (Life Technologies). Relative transcript abundance was determined using the ^ΔΔ^C_T_ method in a SYBR Green assay with GusB and ActinB used in combination for normalisation to housekeeping genes. The following forward (F) and reverse (R) primers were used in mouse samples: Acta2 (F: GTCCCAGACATCAGGGAGTAA and R: TCGGATACTTCAGCGTCAGGA), *Sox9* (F: GGCCGAAGAGGCCACGGAAC and R: GATTGCCCAGAGTGCTCGCCC), *Col1* (F: TGGACGGCTGCACGAGTCAC and R: GCAGGCGGGAGGTCTTGGTG), *Mlc2* (F: CTCTGCAGCAGGGAAACCC and R: CTTCTTGGTGGTCTTGGCCT), *Yap* (F: ATTTCGGCAGGCAATACGGA and R: CATCCTGCTCCAGTGTAGGC), *Itgβ1* (F: GCCAAGTGGGACACGGGTGAA and R: AGCTTGGTGTTGCAAAATCCGCCT), *GusB* (F: GCAGTTGTGTGGGTGAATGG and R: GGGTCAGTGTGTTGTTGATGG), *β-actin* (F: GCTGTATTCCCCTCCATCGTG and R: CACGGTTGGCCTTAGGGTTCAG)

For qPCR in human LX2 cells the following forward (F) and reverse (R) primers were used:

*SETDB1* (F: TAAGACTTGGCACAAAGGCAC and R: TCCCCGACAGTAGACTCTTTC), *SUV3*9*H1* (F: CCTGCCCTCGGTATCTCTAAG and R: ATATCCACGCCATTTCACCAG), *COL1A1* (F: TGTTCAGCTTTGTGGACCTCCG and R: CGCAGGTGATTGGTGGGATGTCT), *ACTA2* (F: CCCCGGGACTAAGACGGGAATC and R: AAGCCGGCCTTACAGAGCCCA), *GUSB* (F: CTCATTTGGAATTTTGCCGATT and R: CCGAGTGAAGATCCCCTTTTT), *β-ACTIN* (F: CCAACCGCGAGAAGATGA and R: CCAGAGGCGTACAGGGATAG)

### Histology, immunohistochemistry and RNAScope

Samples for histology were fixed in 4% PFA overnight followed by storage in 70% ethanol prior to embedding in paraffin and sectioning at 5µm thickness. Histological staining and quantification was performed as previously described^3, 9^. All livers were embedded in an identical orientation and corresponding histology images within the manuscript are shown from the same liver lobe. The extent of scarring was assessed in livers using picro-sirius red (PSR) staining (Sigma). Immunohistochemistry for α-SMA (NCL-L-SMA, Leica Biosystems, dilution: 1:100) and CK19 (ab52625, Abcam, 1:500) was detected using diaminobenzidine (DAB) and Immpact Vector Red (Vector Laboratories) respectively, counterstained with toluidine blue.

*In situ hybridization* for *Pak1* was performed using an RNAscope® 2.5 LS Reagent Kit-RED kit (Cat. No.322150) according to manufacturer’s instructions using a Leica Bond RX automated staining system^9, 43^. After baking and de-waxing slides for 15 minutes, they were pre-treated with HIER for 15 minutes at 95 degrees and 15 minutes of protease digestion. After washing slides with the BOND wash solution (Leica), slides were incubated with *Pak1* RNA probes for 2 hours. Detection of the probes was performed through the BOND polymer Refine Red Detection and haematoxylin kit (DS9390, Leica Bond). Haematoxylin (blue) was used as a counterstain. Slides were heat dried for 30 minutes at 60 degrees. Following a single dip in pure xylene, slides were mounted (EcoMount, Biocare Medical) and cover-slipped.

Quantification of PSR and α-SMA and CK19 staining was determined by morphometric analysis from images acquired using the 3D Histech Pannoramic 250 Flash II slide scanner. From all three lobes of the liver (100 x total magnification), 10 regions were selected at random, and analysed using Adobe Photoshop. The Colour Range tool was used to select stained pixels (red for PSR, brown for α-SMA and red for CK19); the number of selected pixels was recorded and expressed as a fraction of the total number of pixels, averaged across the 10 different regions per section. All quantification was carried out blinded, without prior knowledge of sections or treatment / control group.

### Immunofluorescence, micropatterns and nuclear morphology analyses

Isolated WT and *Pak1*-null HSCs were plated onto chamber slides or hydrogels to activate. Hydrogels of the indicated stiffness (Softwell plates, Cell Guidance Systems) were coated with fibronectin prior to seeding to ensure cell attachment. For micropattern analysis, cells were cultured on plastic for 5 days for activation before trypsinisation and plating onto fibronectin coated micropatterned coverslips (Cytoo), proceeding to fixation 30 minutes after plating to allow adhesion and shape changes. At the indicated time point, media was removed, cells were washed twice in PBS and fixed in 4% PFA in PBS for 10 minutes. After 2 further PBS washes cells were stored under PBS at 4 degrees until staining. Cells were stained with Alexa Fluor 488 phalloidin (Invitrogen 1:500) and/or the respective primary and secondary antibodies in a standard immunofluorescence protocol. Antibodies used were: anti-H3K9Me3 (#13969, CST, 1:500), anti-α-SMA (NCL-L-SMA, Leica, 1:100), anti-SOX9 (ab5535, Merck, 1:1000), anti-YAP (#4912, CST, 1:100), anti-LMNA/C (#2032, CST, 1:500), anti-Vimentin (ab24525, Abcam, 1:500) with their respective species specific Alexa Fluor conjugated secondary antibodies (Invitrogen 1:500). Cells were mounted in Vectashield +DAPI to visualise the nucleus. Images were collected on a Zeiss Axioimager D2 upright microscope and captured using a Coolsnap HQ2 camera (Photometrics) through Micromanager software v1.4.23.

For nuclear morphology analysis, In FIJI images from the DAPI channel were thresholded using the Image>Adjust>Threshold tool using a consistent value across images to define the nuclear area.

These areas were saved as an overlay using the ROI manager. The overlay was applied to the original DAPI image to demarcate the nuclei and the nuclear areas were measured to provide average signal intensity and nuclear morphology data.

### Live F-actin, contraction and cell migration

To visualise live F-actin, WT and *Pak1*-null HSCs were plated at a density of 15,000 cells per well and incubated at 37°C in a glass-bottomed 24-well plate (Cellvis, USA). SiR-actin and Verapamil (Cytoskeleton Inc., USA) were added to cell media at a concentration of 1μl/ml and incubated for a minimum of 6 hours at 37°C. Ten images per animal were taken at x20 magnification using a 3i spinning disc confocal microscope (Leica) and SlideBook 6 software (Intelligent Imaging, USA). Saturated SiR-actin fluorescence was quantified using colour detection analysis in Photoshop software. The background in each image was deducted and saturated SiR-actin staining was calculated as a percentage of the remaining cell area^43^.

To study cell migration, HSCs at 75% confluency were trypsinised and plated into 24 well plates at a density of 5,000 cells per well with 16% growth medium and incubated overnight at 37°C. The next day, plates were loaded onto a wide field microscope with time-lapse capability (Nikon LT TL). NIS-Elements AR software (Nikon) was used to acquire images from predetermined points at x20 magnification every 10 minutes over a 24-hour period. Image sequences were analysed using ImageJ software. The MTrackJ plug-in (ImageJ) was used to track cell movement over the 24-hour period. To determine relative cell migration, six random cells were tracked, and the total cell track distance was measured. For individual cells, the co-ordinates at each ten-minute interval were made relative to a starting point (0,0) and plotted to give representative distance graphs. Measurements in ImageJ were generated in arbitrary units^3, 43^. Directionality was determined by calculating the hypotenuse length of the triangle outlined by the minimum and maximum positions on the x and y axis.

For gel contraction assays, HSCs in culture were detached using trypsin, centrifuged and resuspended in 16% growth medium at a density of 40,000 cells per ml. In a 12-well plate collagen gels were created using 295μl of rat tail collagen (Corning), 6.66μl of newly titrated 1M NaOH and 700μl of HSCs in 16% growth medium per gel. Plates containing the seeded gels were incubated at 37°C for one hour. After solidifying, gels were transferred into a well of a 6-well plate containing 3ml of prewarmed media while avoiding tearing or folding in the gel. Images were taken at 0 hour and 24-hours, following incubation at 37°C, using the colorimetric imaging settings of a transilluminator (Bio-Rad). To measure contraction, ImageJ software (ImageJ) was used to draw a circle around the internal edge of the well and the perimeter of the gel at each time point. Gel sizes were calculated as a percentage of the well area, and contraction calculated as the percentage change between 0 hour and 24 hours.

### Atomic Force Microscopy (AFM)

AFM experiments were performed on a Nanowizard IV AFM (JPK) mounted on an Axio Observer.A1 (Zeiss) inverted optical microscope, with a Vortis Controller (JPK) using SPM Desktop software v6.1.65 (JPK). Isolated WT and *Pak1*-null HSCs were plated onto fibronectin-coated glass dishes for live cell imaging, or fibronectin coated glass coverslips for fixed cell analysis. For live cell analysis, samples were held in a heated petri dish holder during measurements. For fixed cell analysis, cells were fixed in 2.5% glutaraldehyde at the indicated time point and stored under PBS until analysis. Mechanical data was obtained in fluid using 5μm diameter gold colloidal probes (CP-CONT-AU-B, sQUBE, Germany) mounted on cantilevers with a nominal spring constant of 0.3N/m; exact values for cantilever spring constants were determined prior to data acquisition using the JPK non-contact method. Force curves were analysed using JPK Data Analysis software to calculate the average Young’s Modulus of the nucleus based on the Hertz model. When comparing between groups (eg. *Pak1*-null vs WT), measurements were taken with a consistent setpoint and approach velocity for all nuclei analysed and using the exact same probe.

### RNA Microarray

HSCs were isolated from WT and *Pak1*-null mice and RNA was extracted on day 0 (quiescent) or after 10 days (activated) of culturing on plastic. RNA was extracted as previously described and stored at -80°. The integrity of RNA samples was assessed, and samples were considered of sufficient quality when having an RNA integrity number (RINe) of ≥6.5. An Affymetrix mouse genome 430 2.0 array using 2894 probe sets was employed for this experiment. Data was initially assessed using principal component analysis (PCA) to determine the source of variation in results, before undergoing cluster analysis. Subsequent datasets were then examined using online bioinformatic analytical software. The Database for Annotation, Visualization and Integrated Discovery (DAVID) analysis version 6.8 (https://david.ncifcrf.gov/) was used for functional annotation and gene ontology analysis, whilst Ingenuity Pathway Analysis (https://www.qiagenbioinformatics.com) was employed to identify key canonical pathways between experimental groups.

### Cut&Tag

Cut&Tag was performed using a Complete CUT&Tag-IT Assay Kit (Active Motif) according to manufacturer’s instructions. For each reaction of quiescent samples, 500,000 cells were counted from a pool of freshly isolated HSCs from three mice. For activated samples, mouse HSCs were cultured on plastic for 7 days and collected by brief incubation at 37 degrees in 0.05M EDTA in HBSS to promote detachment followed by cell scraping. For each activated reaction, 500,000 cells were counted from a pool of HSCs from three mice. Briefly, cells were attached to concanavalin A beads before overnight binding of primary antibody. ‘H3K9Me3 antibody 1’ refers to 39765 (Active Motif), ‘H3K9Me3 antibody 2’ refers to 13969 (CST) and the H3K27Me3 to 39155 (Active Motif). This preceded binding of secondary antibody and CUT&Tag-IT™Assembled pA-Tn5 Transposomes, tagmentation and purification of tagmented DNA. This DNA was PCR amplified using a unique combination of i5/i7 indexing primers for each sample, cleaned using SPRI beads and pooled into an equimolar library for sequencing on an Illumina HiSeq4000 sequencing platform.

Unmapped paired-end reads of 76bp were checked using a quality control pipeline consisting of FastQC v0.11.3 (http://www.bioinformatics.babraham.ac.uk/projects/fastqc/) and FastQ Screen v0.14.0 (https://www.bioinformatics.babraham.ac.uk/projects/fastq_screen/). Reads were trimmed to remove any remaining adapters and poor quality sequence using Trimmomatic v0.39^44^; reads were truncated at a sliding 4bp window, starting 5’, with a mean quality <Q20, and removed if the final length was less than 35bp. Additional flags included: ‘ILLUMINACLIP:./Truseq3-PE-2_Nextera-PE.fa:2:30:10 SLIDINGWINDOW:4:20 MINLEN:35’.

Paired-reads were mapped to the mouse genome (mm39/GRCm39)^45^ using Bowtie2 v2.4.2^46^. samtools v1.12^47^ was used to create (view) and sort (sort) BAM files from the SAM files output from Bowtie2. Reads located in blacklist regions were removed using bedtools intersect v2.27.1^48^. Blacklist regions for mouse mm10 (https://github.com/Boyle-Lab/Blacklist/blob/master/lists/mm10-blacklist.v2.bed.gz) were converted to mm39 using UCSC liftOver, the ‘Minimum ratio of bases that must remap’ was set to 0.95. The reads were then filtered to retain only concordant read pairs with a minimum quality score of 2, using samtools view (-f2 -q2).

Prior to peak calling read pairs were removed that mapped to the mitochondrial genome or unassembled contigs, using the Linux bash tool ‘sed’ (sed ‘/chrM/d;/random/d;/chrUn/d;/GL456/d;/JH584/d;/MU069/d’). Candidate binding regions ‘peaks’ were identified using SEACR v1.3^49^. The software was run in both ‘relaxed’ and ‘stringent’ modes. In both sets of fragment counts were normalised ‘norm’, and the top 1% of regions were selected. The closest gene was associated with each region using RnaChipIntegrator v2.0.0 (https://github.com/fls-bioinformatics-core/RnaChipIntegrator) (--cutoff=100000 --edge=both --number=1 --compact). The gene annotation for Gencode vM27 / mm39 (knownCanonical) was downloaded using the UCSC table browser.

### ATAC-seq/RNA-seq

Accessible genomic DNA was obtained for sequencing using an ATAC-seq kit (Active Motif) as per manufacturer’s instructions. 100,000 freshly isolated mHSCs were used for each quiescent reaction. For activated samples, mHSCs were cultured on plastic for 7 days and collected by brief incubation at 37 degrees in 0.05M EDTA in HBSS to promote detachment followed by cell scraping, and 100,000 cells used for each reaction. Cells were pelleted, washed with PBS then resuspended in ATAC lysis buffer. Nuclei were pelleted by centrifugation at 500g at 4 degrees before resuspension in tagmentation buffer and incubation at 37 degrees for 30 minutes. DNA was then purified and each sample amplified with a unique combination of i5/i7 primers for 10 cycles followed by SPRI clean-up and pooling into an equimolar library. In parallel, RNA was extracted from quiescent and activated cells using an RNeasy kit (Qiagen) for RNA-seq.

For ATAC-seq, unmapped paired-reads of 76bp from an Illumina HiSeq4000 sequencer were QC checked as described for Cut&Tag and removed if the final length was less than 25bp. Paired-reads were mapped to the mouse genome as described for Cut&Tag, using additional parameters (’-X 2000 --very-sensitive’). Picard v2.1.0 MarkDuplicates (https://broadinstitute.github.io/picard/) was used to remove duplicate reads on the same strand. Mitochondrial reads and reads located in blacklist regions were removed as described for Cut&Tag. ATAC-seq fragment length quality plots were generated using ATACseqQC v1.12.5^50^ in R v4.0.2 (R Core Team 2021). Candidate regions of open chromatin ‘peaks’ were identified using MACS2 v2.2.7.1^51^ using additional parameters (--format BAMPE -- gsize mm --keep-dup all --qvalue 0.01 --bdg --SPMR --call-summits). DNA fragment coverage profiles generated by MACS2 were converted into bigWig and bigBed format using the UCSC tools bedClip, bedGraphToBigWig, and bedToBigBed.

Differential binding analysis was performed using DiffBind v3.4.11^52^ in R v4.1.2. The ‘peak’ input consisted of 200bp regions centred on MACS2 summit coordinates, in BED format, together with the qvalue of each region (chr, start, end, q-value). The ‘read’ input were the final filtered BAM files used in the MACS2 peak calling. dba.count parameters were set differently from the default (minOverlap=2, summits=FALSE).

UCSC knownCanonical genes from Gencode Mouse v27 were associated with Diffbind regions using RnaChipIntegrator v2.0.0, including the following parameters (--cutoff=100000 --edge=both -- number=2 --compact). In order to identify DiffBind regions in the promoters of genes the following parameters were used (--cutoff=100000 --edge=tss --number=1 --compact).

For RNA-seq, unmapped paired-reads of 51bp from a NovaSeq6000 were QC checked and trimmed as previously and removed if the final length was less than 35bp. Additional flags included: ‘ILLUMINACLIP:./Truseq3-PE-2_Nextera-PE.fa:2:30:10’. The filtered reads were mapped to the mouse reference sequence analysis set (mm39/GRCm39) from the UCSC browser (Kent et al. 2002), using STAR v2.7.7a (Dobin et al. 2013). The genome index was created using the comprehensive mouse Gencode v27 gene annotation (Harrow et al. 2012) applying a read overhang (--sjdbOverhang 75). During mapping the flags ‘--quantMode GeneCounts’ was used to generate read counts into genes. Normalisation and differential expression analysis was performed using DESeq2 v1.34.0 on R v4.1.2 (Love, Huber, and Anders 2014). A single factor was included in the model to compare activated versus quiescent cells. Log fold change shrinkage was applied using the lfcShrink function along with the “apeglm” algorithm.

EnrichR was used to identify significantly enriched GO terms within filtered gene lists^53^. Simple Enrichment Analysis from the MEME suite toolset was used to identify enriched TF motifs within promotor ATAC peaks^54^.

### Statistical analysis

Statistical analyses were performed using GraphPad Prism 8 software using the indicated tests. Shapiro-Wilks tests were used to assess normality of data. Where data were normally distributed, two-tailed student t-tests were used for pairwise analyses and one-way ANOVA with Tukey’s post hoc test for comparisons across more than two groups. For non-normally distributed data, Mann-Whitney testing was used for pairwise analysis. Correlation analysis was assessed using a 2-parameter exponential regression model (R-squared).

## Contributions

KPH conceived and designed experiments. EJ, AFM, KM, NWH, CJW and NAH contributed to experimental design. EJ, AFM, KS, LB, LP, KM, JP, NWH, CJW and VSA performed experiments. VSA and RS provided human tissues. EJ, AFM, EC, LZ and KPH analyzed the data. LZ and ID performed bioinformatics. EC carried out complex statistics. KPH and EJ guided experiments and wrote the manuscript.

## Acknowledgements

This work was supported by the Medical Research Council (MRC; KPH, MR/J003352/1 & MR/P023541/1; NAH, MR/000638/1 & MR/S036121/1; and KM and VSA are MRC Clinical Training Fellows). We thank Sara Wickstrom and her team at the University of Helsinki for technical and experimental guidance. The Genomics Core Facility and the Bioimaging Facility at the University of Manchester are acknowledged for their technical help and support from Wellcome (105610). KPH is a member of the Wellcome Trust supported Centre for Cell-Matrix Research (203128/Z/16/Z). This report is independent research supported by the North West Lung Centre Charity at Manchester University NHS Foundation Trust. The authors would like to acknowledge the Manchester Allergy, Respiratory and Thoracic Surgery Biobank and the North West Lung Centre Charity for supporting this project. The views expressed in this publication are those of the author(s) and not necessarily those of the NHS, the North West Lung Centre Charity or the Department of Health.

