## Supplementary for "PAK1-dependant mechanotransduction enables myofibroblast nuclear adaptation and chromatin organisation during fibrosis"

***This document includes Supplementary Methods, References, Figures 1-13 and Tables 1-2. Supplementary Data sets 1-3 are available as separate Excel files.**

#### Supplementary Methods

#### Primary cell isolation and culture

**qPCR primer sequences**

| Gene |  | Primer sequence (5’-3’) | Species |
| --- | --- | --- | --- |
| *Acta2* | F | GTCCCAGACATCAGGGAGTAA | Mouse |
|  | R | TCGGATACTTCAGCGTCAGGA |  |
| *Sox9* | F | GGCCGAAGAGGCCACGGAAC |  |
|  | R | GATTGCCCAGAGTGCTCGCCC |  |
| *Col1* | F | TGGACGGCTGCACGAGTCAC |  |
|  | R | GCAGGCGGGAGGTCTTGGTG |  |
| *Mlc2* | F | CTCTGCAGCAGGGAAACCC |  |
|  | R | CTTCTTGGTGGTCTTGGCCT |  |
| *Yap* | F | ATTTCGGCAGGCAATACGGA |  |
|  | R | CATCCTGCTCCAGTGTAGGC |  |
| *Itgβ1* | F | GCCAAGTGGGACACGGGTGAA |  |
|  | R | AGCTTGGTGTTGCAAAATCCGCCT |  |
| *GusB* | F | GCAGTTGTGTGGGTGAATGG |  |
|  | R | GGGTCAGTGTGTTGTTGATGG |  |
| *β-actin* | F | GCTGTATTCCCCTCCATCGTG |  |
|  | R | CACGGTTGGCCTTAGGGTTCAG |  |
| *SETDB1* | F | TAAGACTTGGCACAAAGGCAC | Human |
|  | R | TCCCCGACAGTAGACTCTTTC |  |
| *SUV39H1* | F | CCTGCCCTCGGTATCTCTAAG |  |
|  | R | ATATCCACGCCATTTCACCAG |  |
| *COL1A1* | F | TGTTCAGCTTTGTGGACCTCCG |  |
|  | R | CGCAGGTGATTGGTGGGATGTCT |  |
| *ACTA2* | F | CCCCGGGACTAAGACGGGAATC |  |
|  | R | AAGCCGGCCTTACAGAGCCCA |  |
| *GUSB* | F | CTCATTTGGAATTTTGCCGATT |  |
|  | R | CCGAGTGAAGATCCCCTTTTT |  |
| *β-ACTIN* | F | CCAACCGCGAGAAGATGA |  |
|  | R | CCAGAGGCGTACAGGGATAG |  |

#### Supplementary Figures

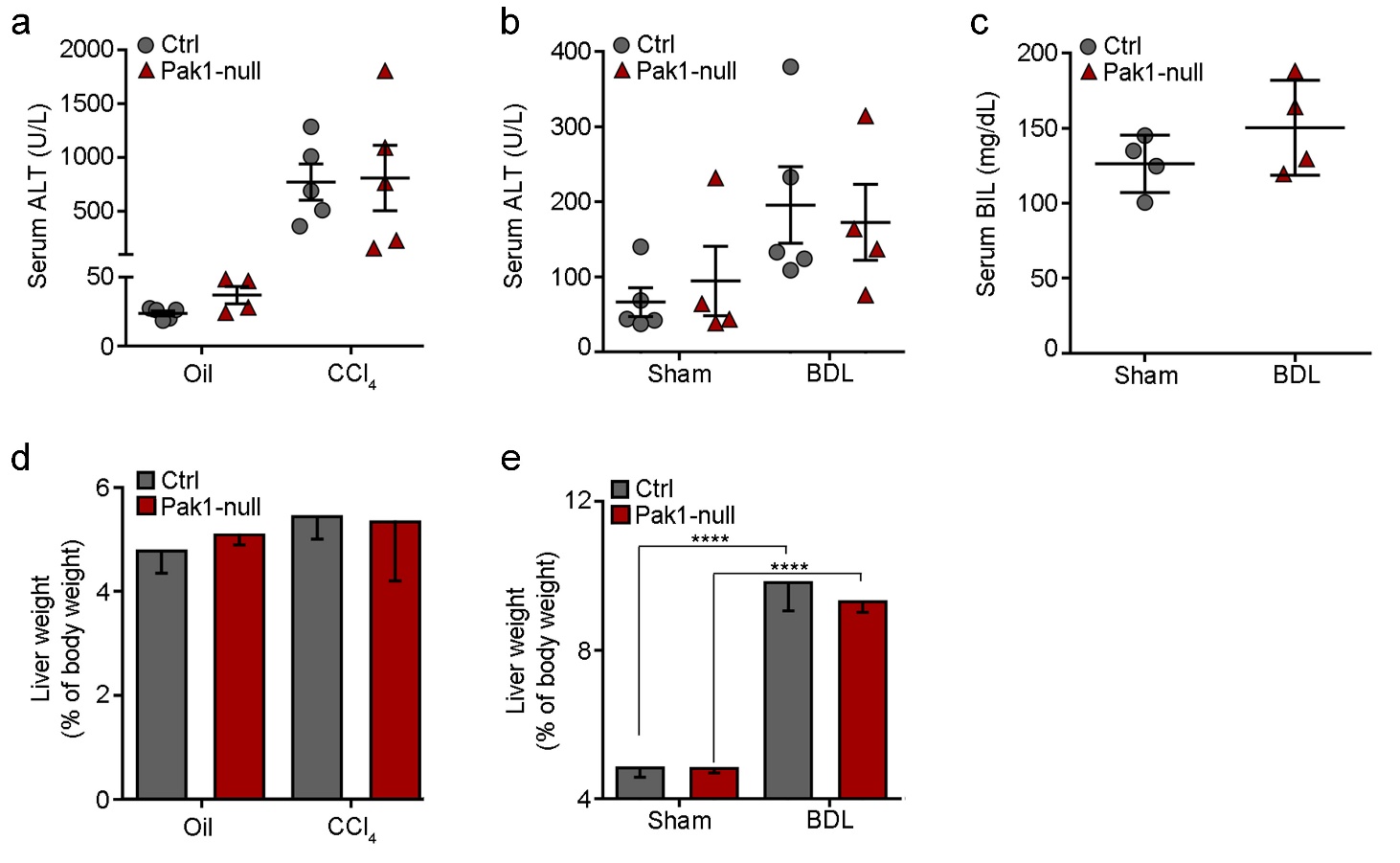
**Supplementary Figure 1** Characterisation of control and Pak1-null livers in CCl_4_ and BDL models of liver fibrosis. (**a-c**) No alterations in liver function in response to Pak1 loss compared to control for serum ALT (CCl_4_ model; **a**) and ALT & bilirubin (BDL; **b, c**). (**d, e**) Pak1 loss did not alter the liver weight/body weight ratio compared to the control group in CCl_4_ (**d**) or BDL (**e**) models of liver fibrosis. Two-tailed unpaired *t*-test was used for statistical analysis. Data are shown as means ± s.e.m. ****P<0.001.

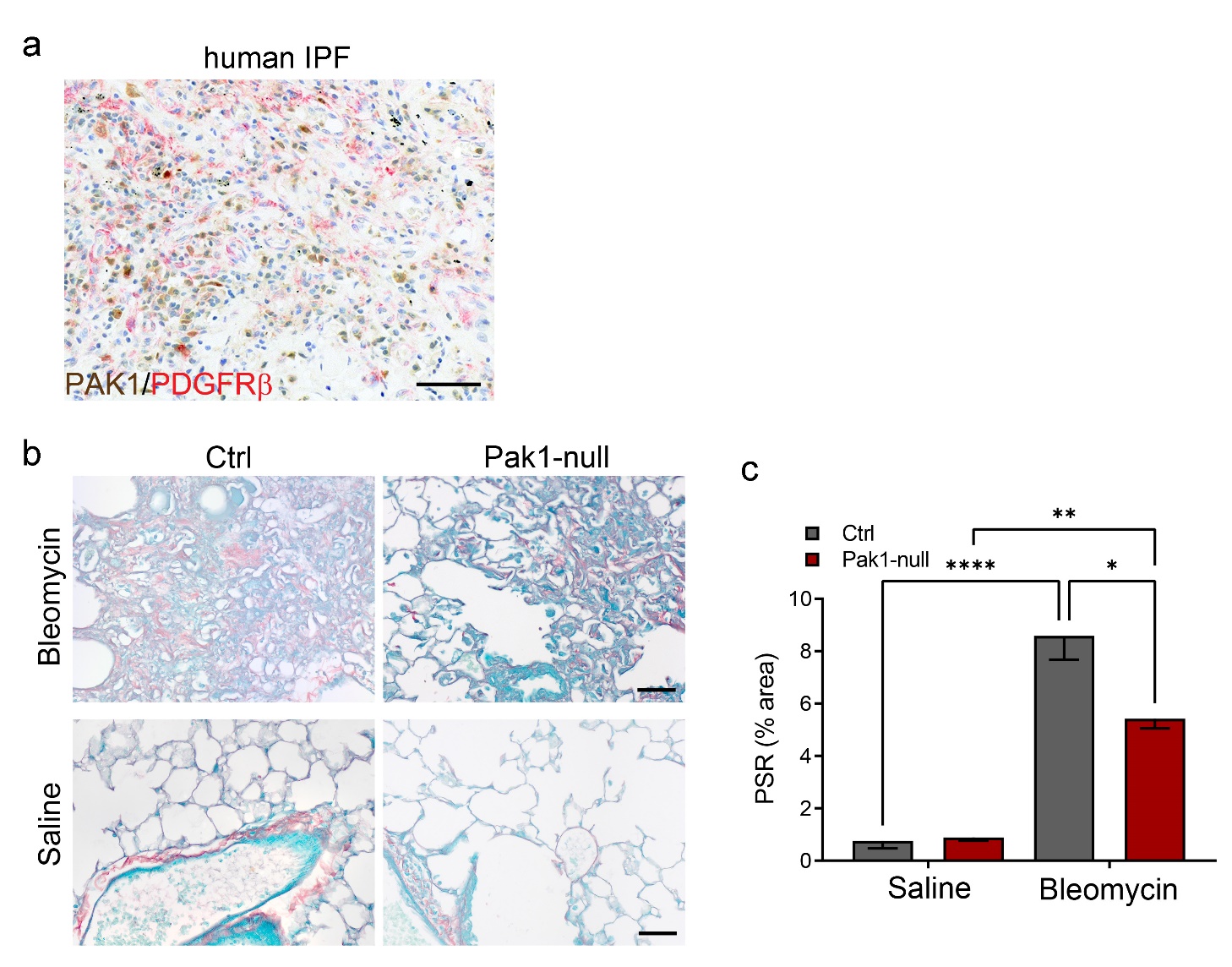

**Supplementary Figure 2** Characterisation of Pak1 in models of human IPF and bleomycin-induced lung fibrosis. (**a**) PAK1 protein (brown) localized to the scar demarcated by PDGFRβ (red) in human IPF tissue. (**b**) PSR lung staining of bleomycin treated Pak1-null and control animals alongside saline controls (n=3 per group), quantified in (**c**). Data are shown as means ± s.e.m. *P<0.05, **P<0.01, ****P<0.001. Two-way ANOVA with Tukey’s multiple comparison test. Size bar = 50 µm.

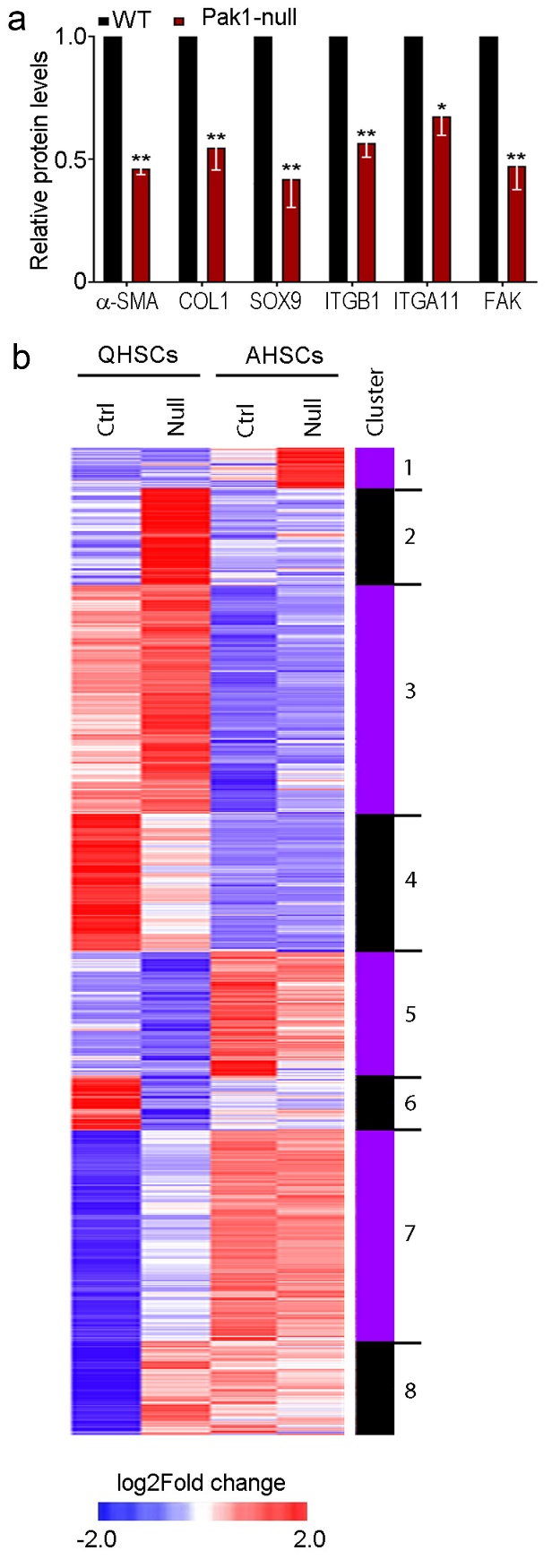

**Supplementary figure 3. PAK1 microarray analysis.** (**a**) Protein quantification by western blot of profibrotic markers in control (wild type, WT) and Pak1-null activated HSCs (n=3 per group). (**b**) Cluster analysis and heatmap of mean gene expression changes (P<0.05) in control (Ctrl) and Pak1-null (Null) quiescent (Q) and activated (A) HSCs. Eight clusters were identified on the basis of increased (red) and decreased (blue) gene expression. Data are shown as means ± s.e.m. *P<0.05, **P<0.01 by two-tailed t-test.

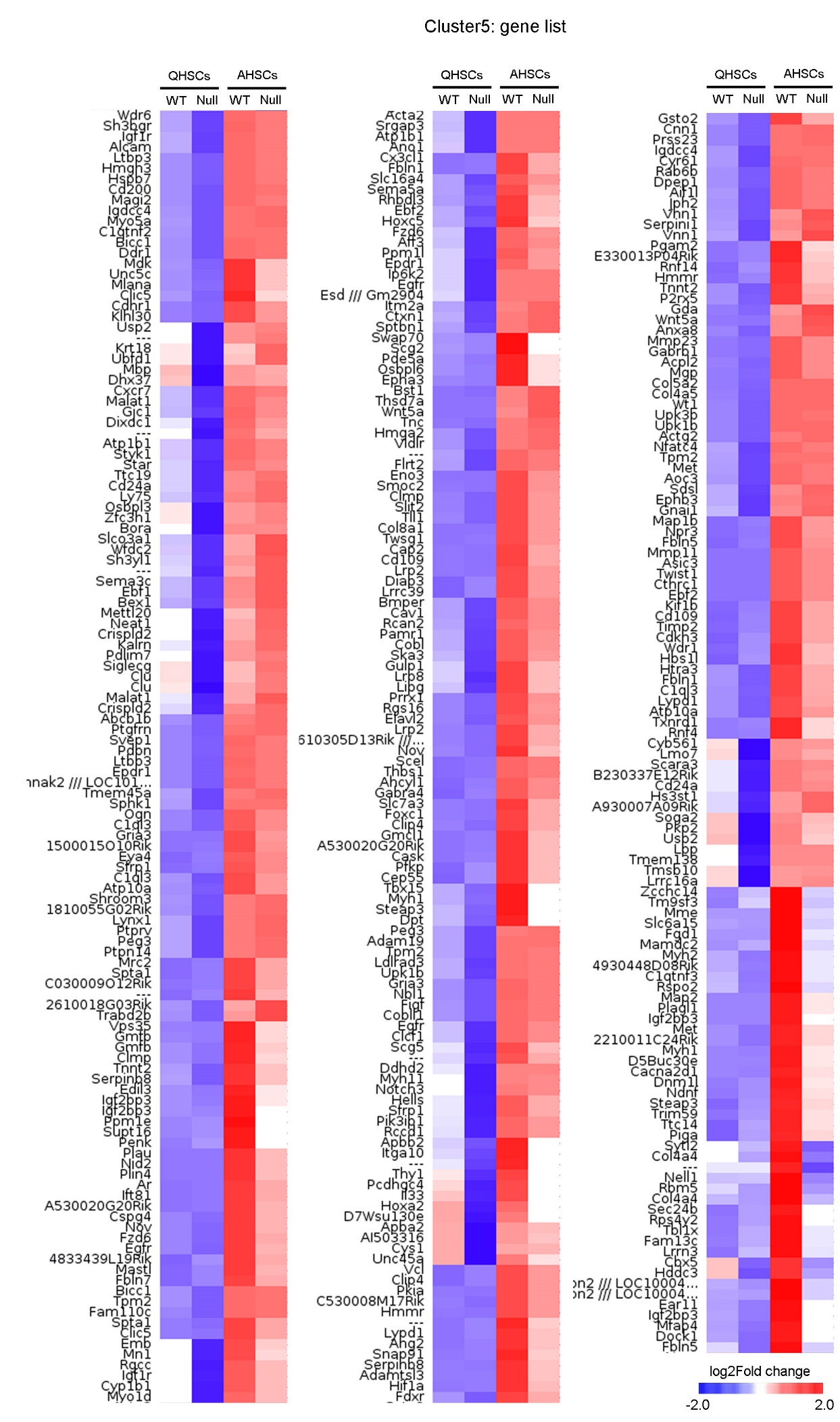
**Supplementary Figure 4** Hierarchical clustering and heatmap for Cluster 5 (Fig. 2b) with full gene list. Upregulated (red) and downregulated (blue) in control (Ctrl) and Pak1-null (Null) quiescent (Q) and activated (A) HSCs.

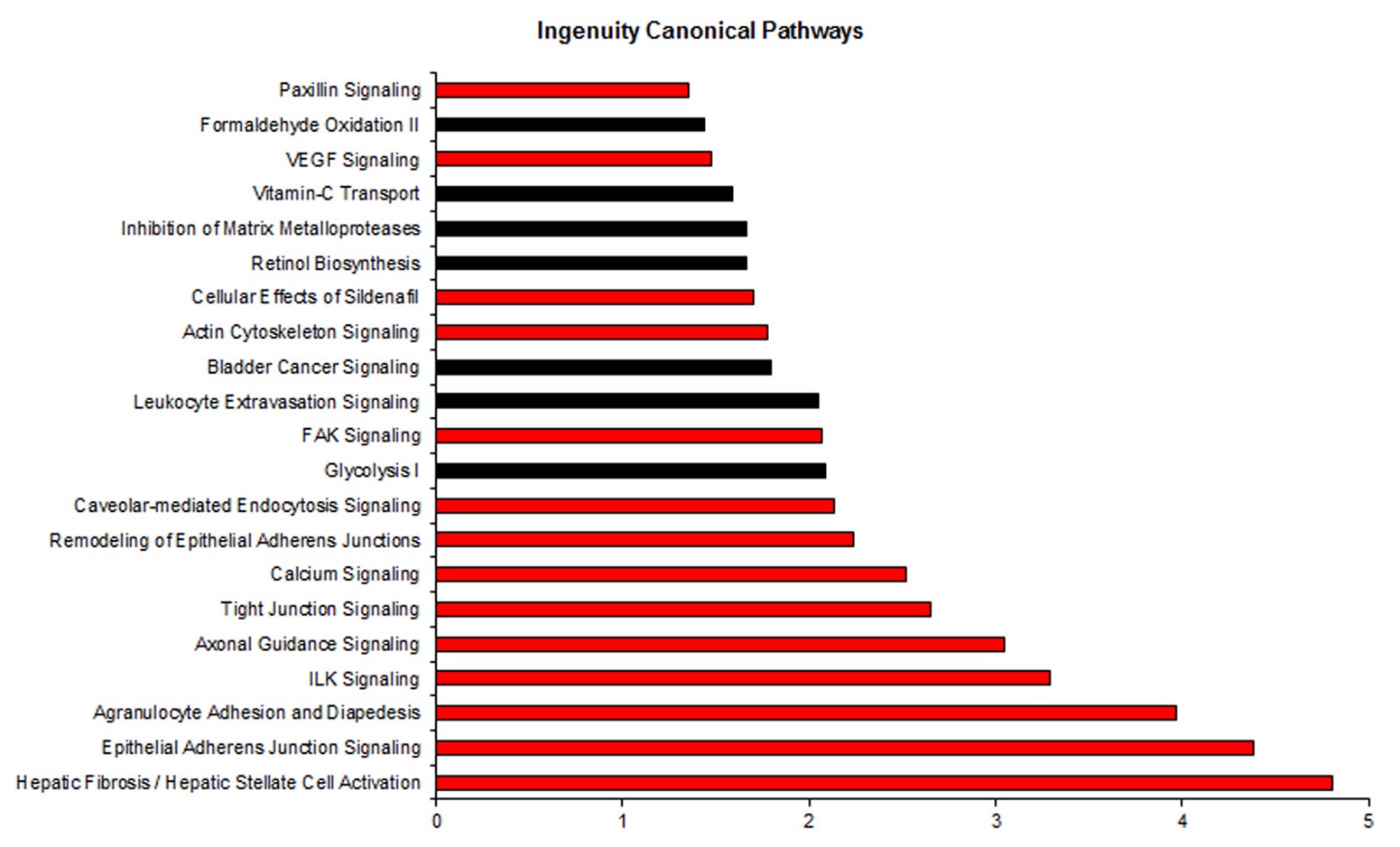
**Supplementary Figure 5** Top canonical pathways represented by genes listed in Cluster 5 (Fig. 2b) following Ingenuity Pathway Analysis. Pathways were ranked by the negative log of P-values calculated by Fisher’s exact test for gene enrichment. Pathways highlighted in red contain either Mechanosignalling as terms.

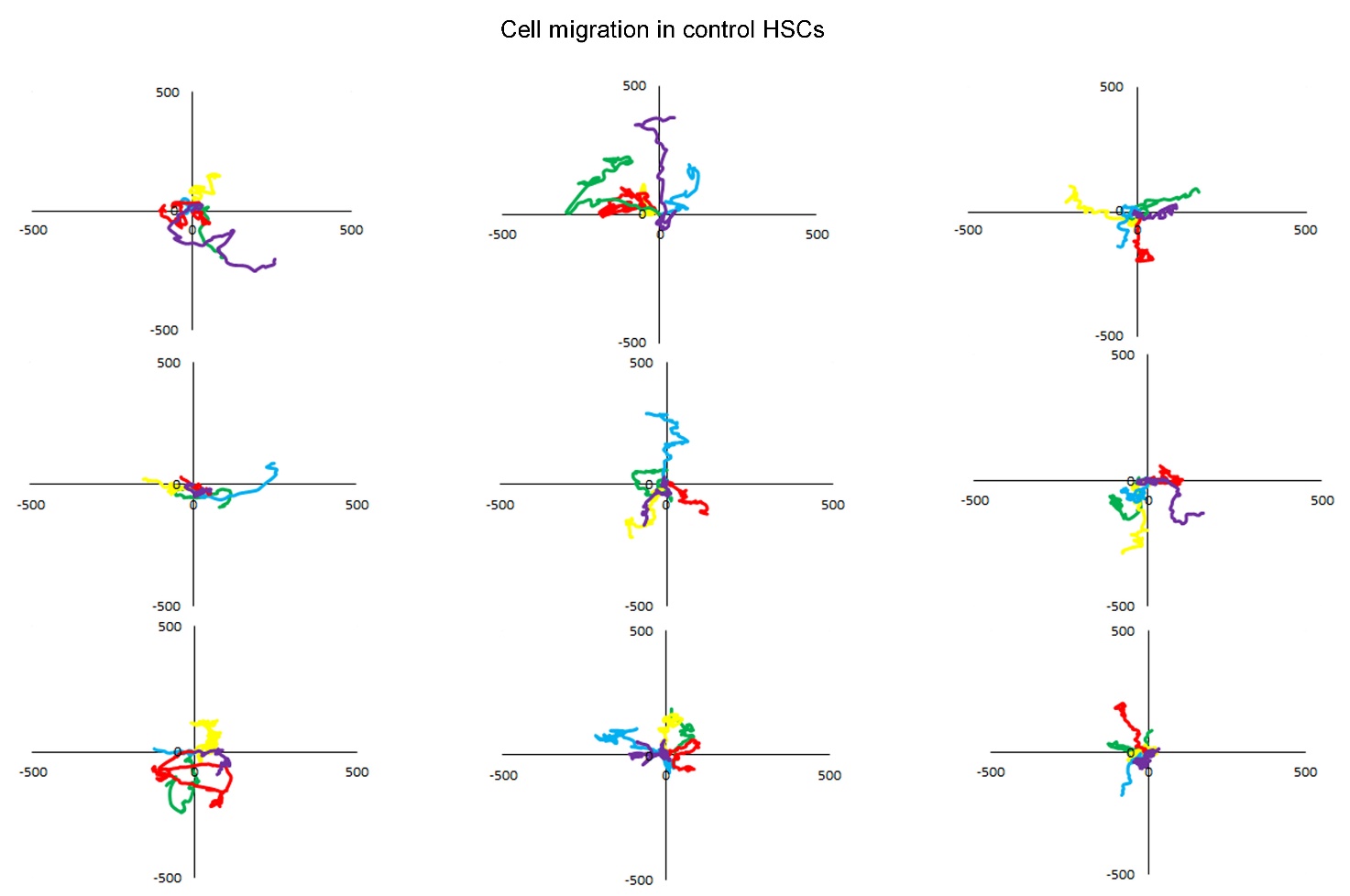

**Supplementary Figure 6** Live cell migration over 24 hours from control AHSCs (shown in Fig 3f). Track length shown for three biological replicates (top, middle and bottom row) and 5 cell tracks for each graph (15 cell tracks for each replicate in total).

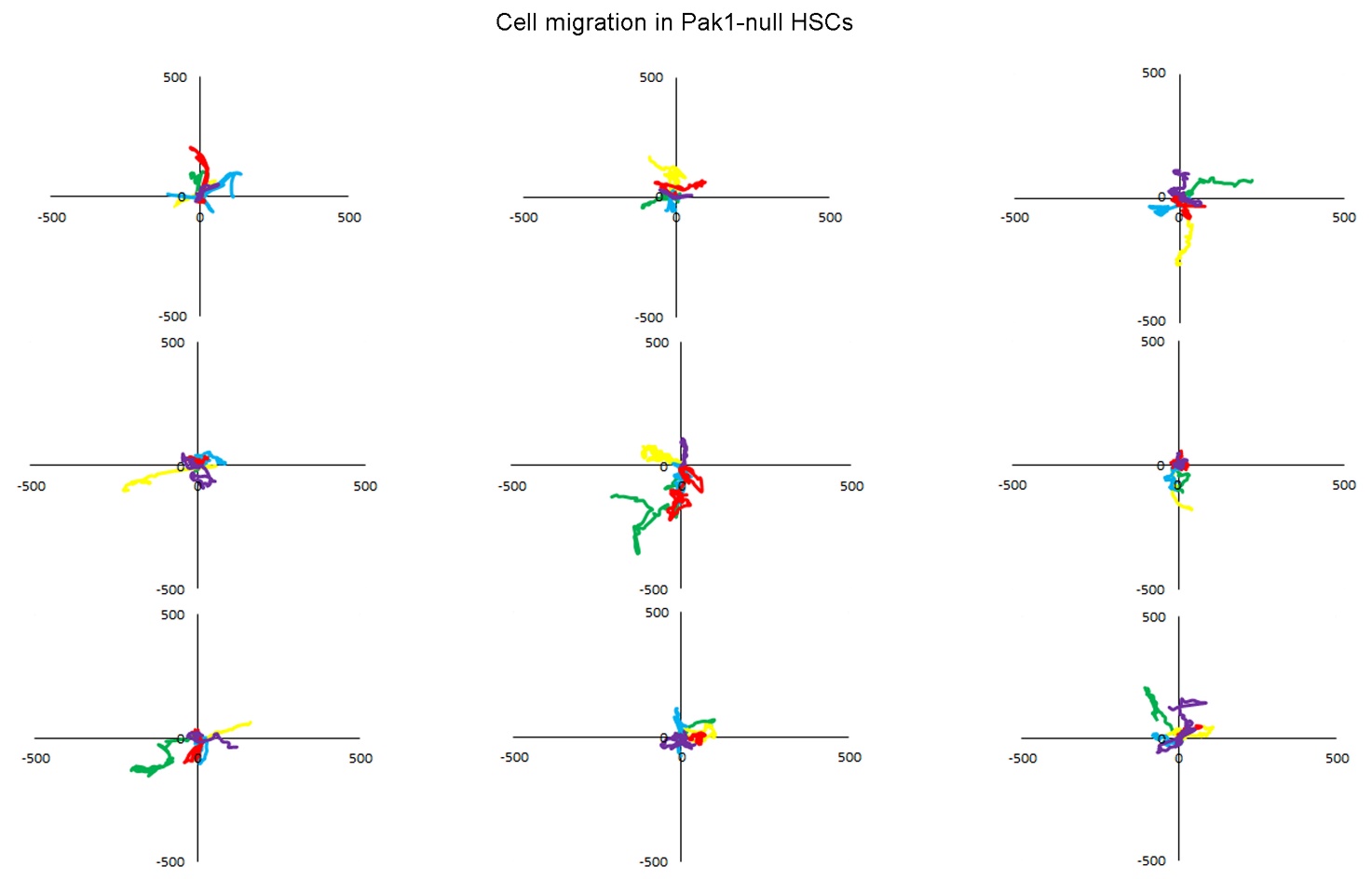

**Supplementary Figure 7** Live cell migration over 24 hours from Pak1-null AHSCs (shown in Fig 3g). Track length shown for three biological replicates (top, middle and bottom row) and 5 cell tracks for each graph (15 cell tracks for each replicate in total).

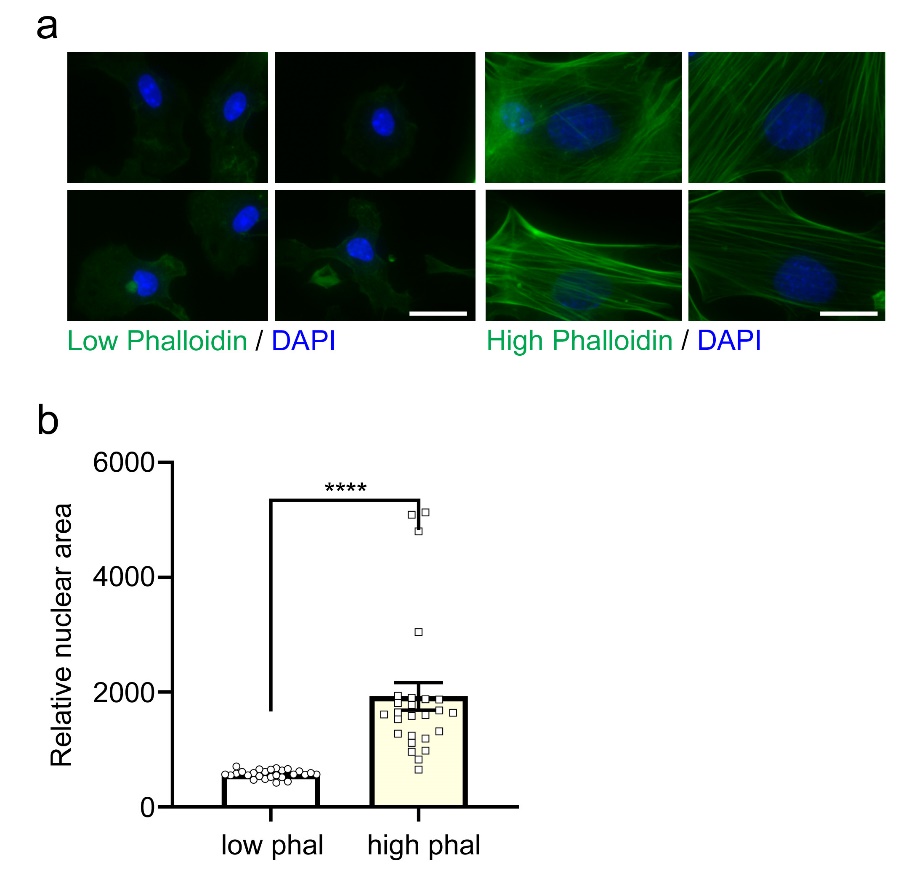

**Supplementary Figure 8** Altered cell state during HSC activation. (**a, b**) Representative DAPI (blue) and Phalloidin (green) images demonstrating activation of HSCs. Altered shape and nuclear area of low phaliodin (corresponding to inactive HSCs) and high phalloidin (corresponding to activated HSCs) are shown in **a** and quantified in **b**. Data are shown as means ± s.e.m. ****P<0.001 using two tailed t-test. Size bar = 10 µm.

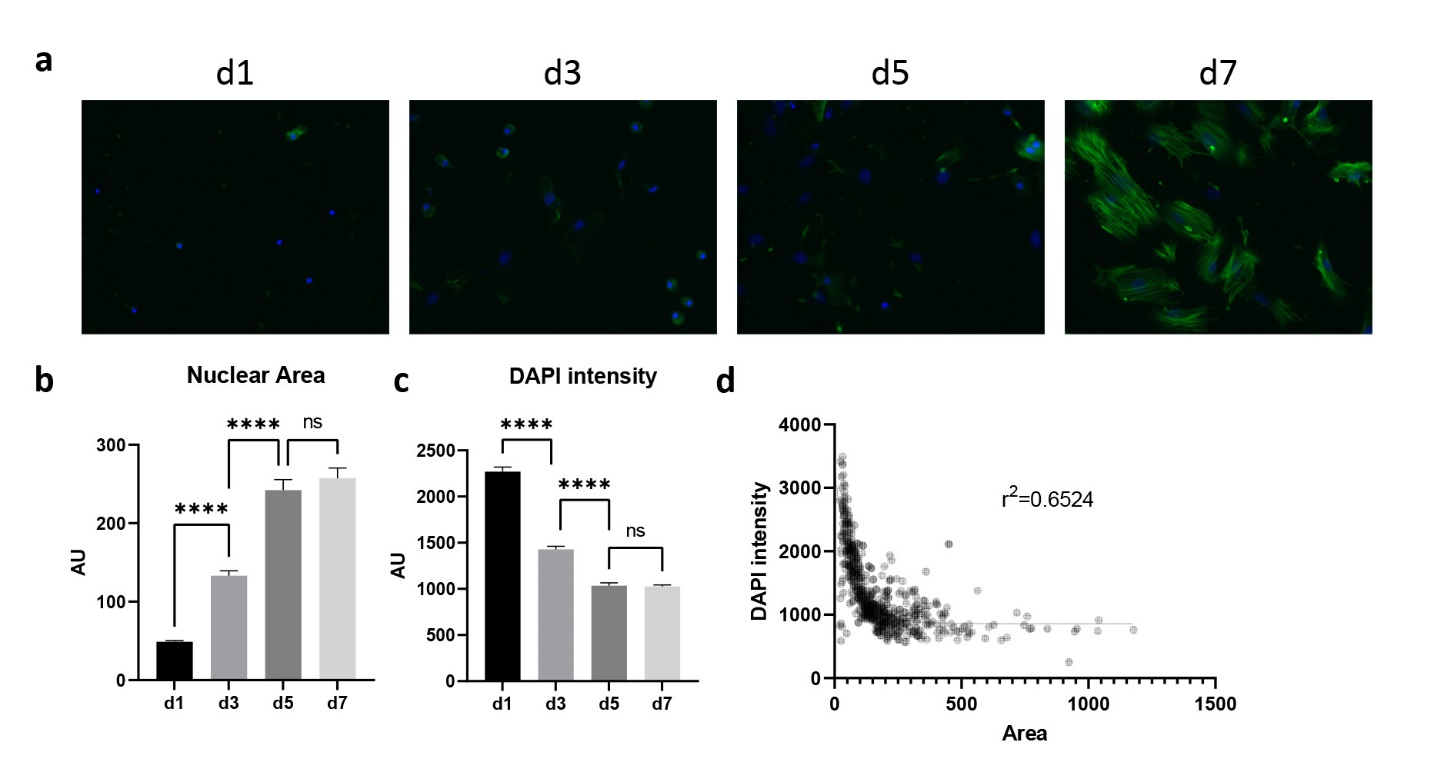

**Supplementary Figure 9** Nuclear expansion of primary lung fibroblasts. (**a**) Activation timecourse of primary lung fibroblasts on plastic, showing an increase in phalloidin staining (green) correlating with nuclear expansion and loss of DAPI (blue) intensity. (**b**) Quantification of nuclear area shows a progressive, significant increase up to day 5 (p<0.001, Kruskal-Wallis test, n>90 cells per time point across three independent mouse preps). (**c**) Quantification of DAPI intensity shows progressive significant decrease up to day 5 (p<0.001, Kruskal-Wallis test, n>90 cells per time point across three independent mouse preps). (**d**) Inverse correlation of DAPI intensity and nuclear area.

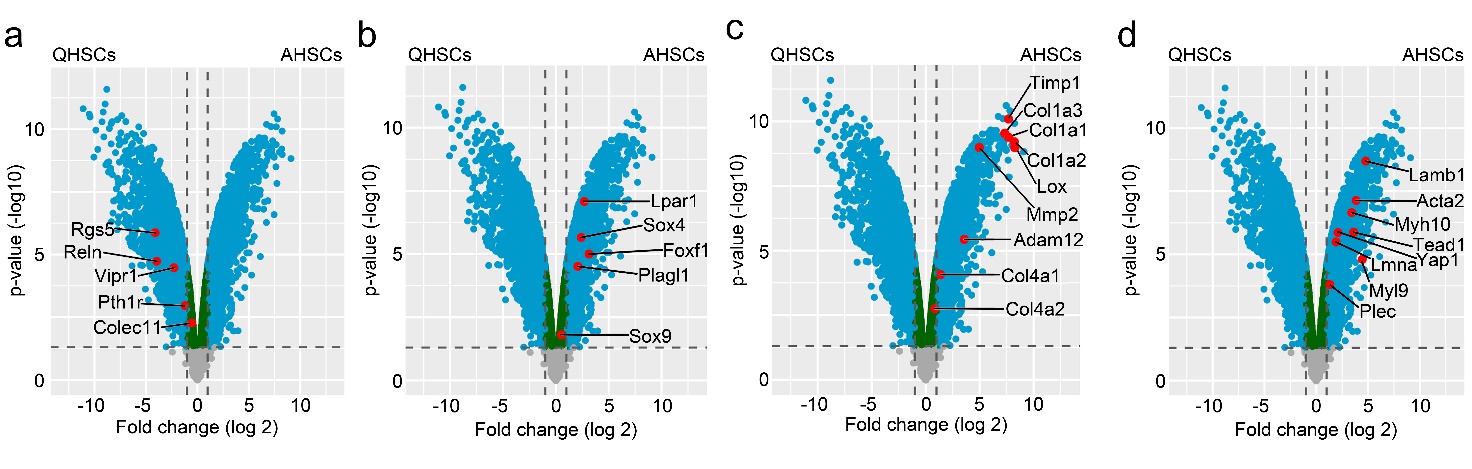

**Supplementary figure 10** (**a-d**) Volcano plots of gene expression changes between quiescent in **a** and activated HSCs in **c-d**. Blue dots, all genes with fold change >2, P<0.05; red dots, highlighted genes representing quiescent (**a**), activated (**b-d**), ECM (**c**) and mechanotransduction (**d**); green dots, genes with fold change < 2, P<0.05; grey dots, unsignificant gene change.

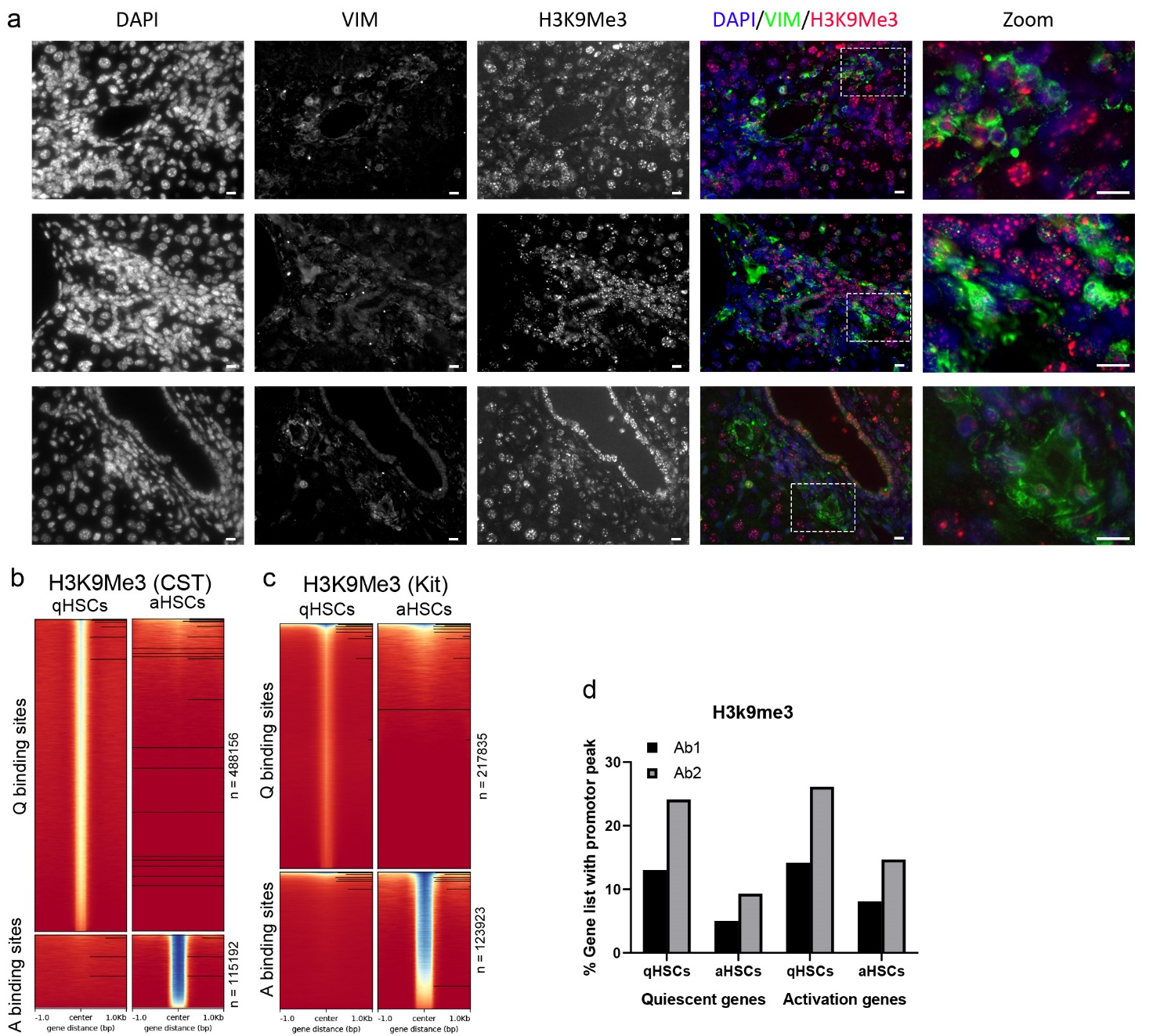

**Supplementary figure 11** (**a**) Extended view of Vimentin (green) and H3K9me3 (red) co-staining showing reduced H3K9me3 in vimentin positive cells across three bile duct ligated animals. (**b-c**) Heatmaps of Cut&Tag binding sites for two independent antibodies against H3k9Me3. n shows the number of peaks in each state. (**d**) Proportion of quiescent genes (>2fold enriched in qHSCs in RNA-seq) and activation genes (likewise for aHSCs) with a Cut&Tag binding site detected on the gene promotor for H3K9me3. Similar reduction during activation is seen in genes known to be up- or downregulated.

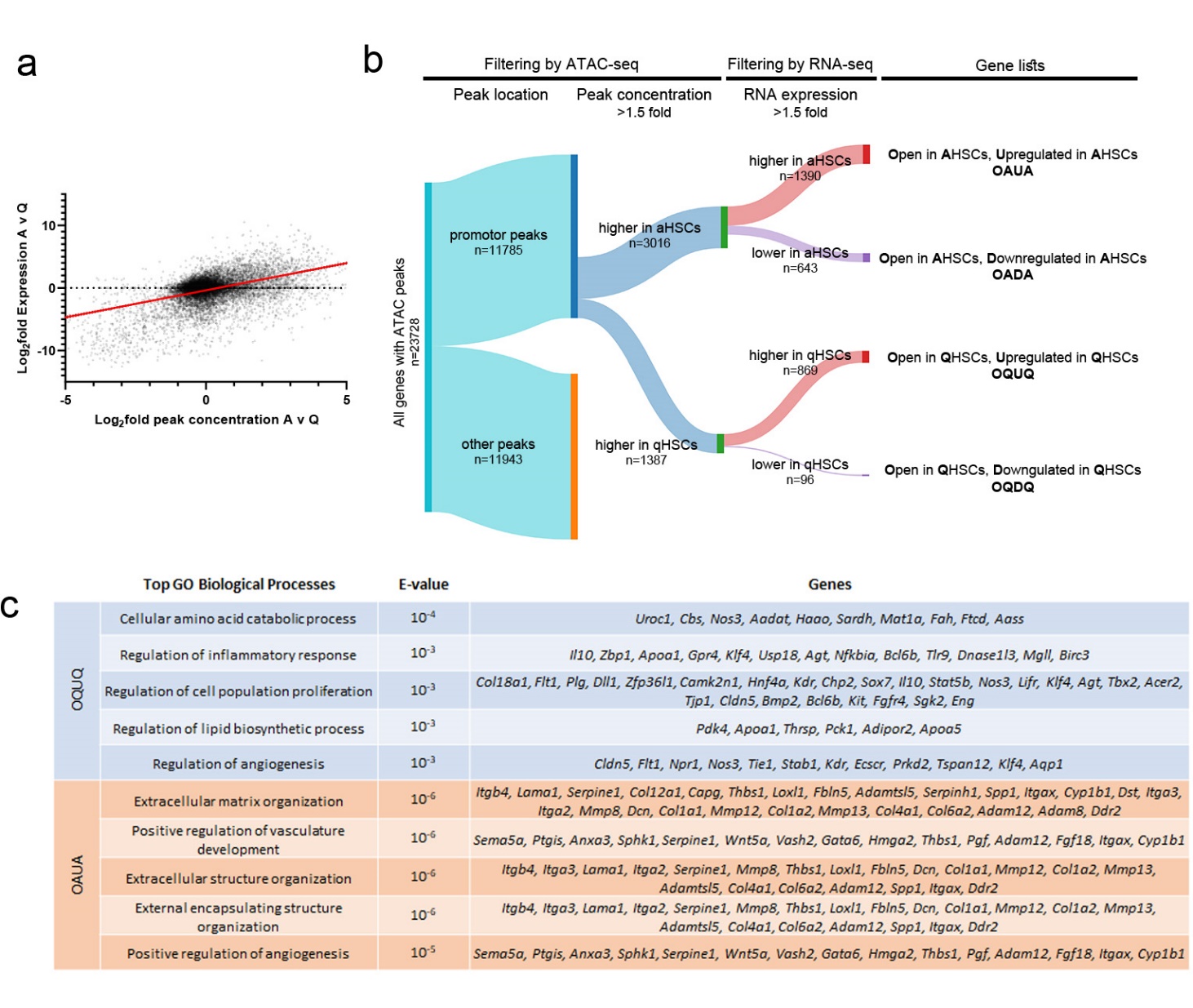

**Supplementary figure 12** (**a**) Correlation of ATAC-seq promotor peak concentration and RNA-seq expression shows a significant positive correlation (Spearman’s rank r=0.328, p<0.0001). (**b**) Sankey diagram showing serial filtering of genes with ATAC peaks overlapping gene promotors based on ATAC-seq promotor peak concentration in quiescent HSCs (Q) or activated HSCs (A). 1.5 fold upregulated in either Q or A and RNA-seq expression 1.5 fold upregulated in either Q or A produced 4 gene lists: chromatin more open and upregulated in A (OAUA), more open and downregulated in A (OADA), more open and upregulated in Q (OQUQ) and more open and downregulated in Q (OQDQ). (**c**) Top 5 GO Biological Processes enriched in genes more open and upregulated in quiescence (OQUQ) and more open and upregulated in activation (OAUA).

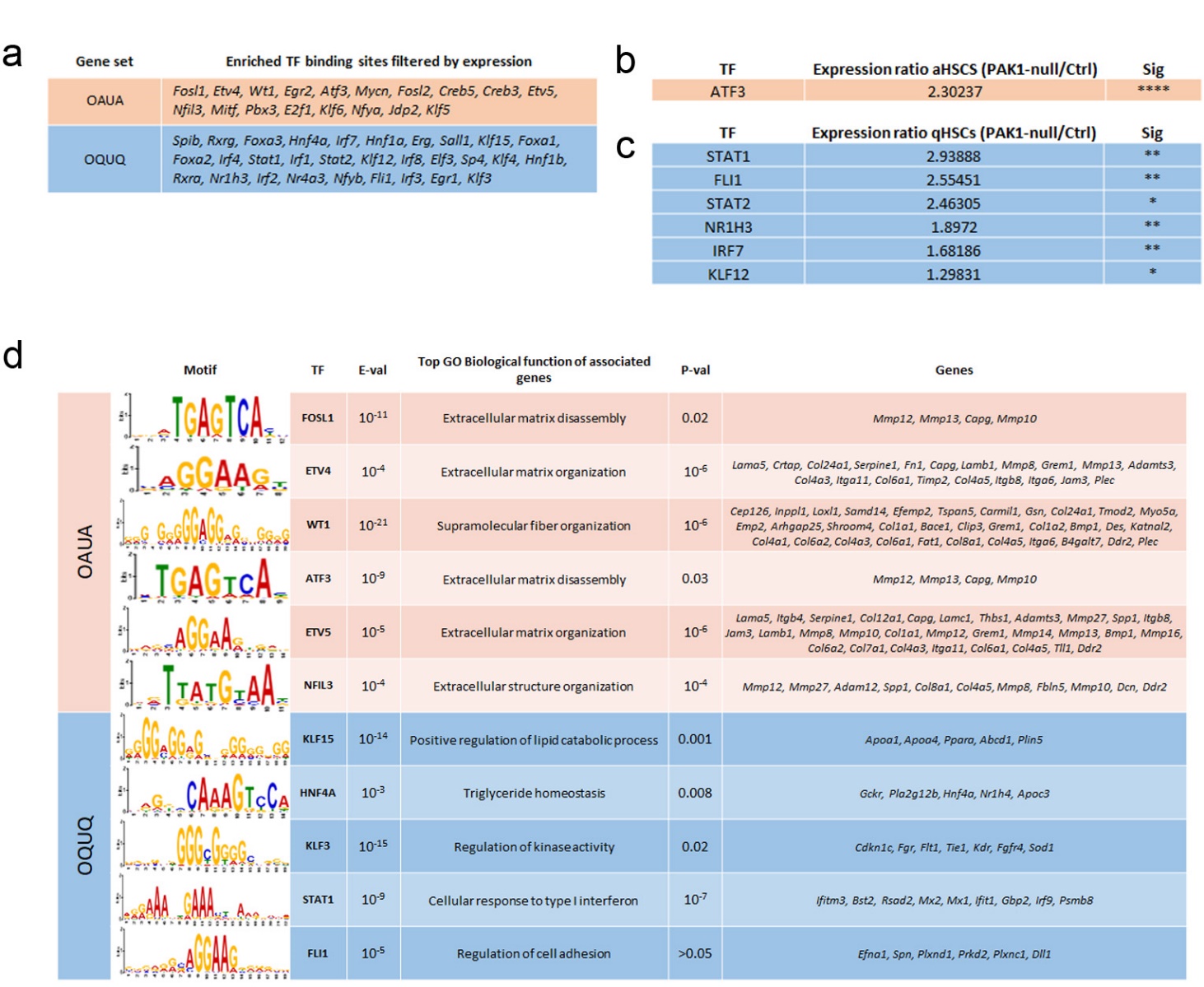

**Supplementary figure 13** (**a**) HSC state-specific TF motifs identified by Simple Enrichment Analysis and filtered to ensure robust expression in RNAseq (>100 reads) and >1.5 fold enrichment in expression in the aHSCs (for OAUA) or qHSCs (for OQUQ). (**b**) Activation-associated TFs with significantly altered expression in PAK1-null aHSCs (**** = p<0.0001). (**c**) Quiescence-associated TFs with significantly altered expression in PAK1-null qHSCs (* = p>0.05, ** = p<0.01). (**d**) Selected HSC state-specific TF motifs and the top GO Biological function of the list of genes containing that motif in the promotor, alongside the list of genes contributing to the GO.

#### Supplementary Tables

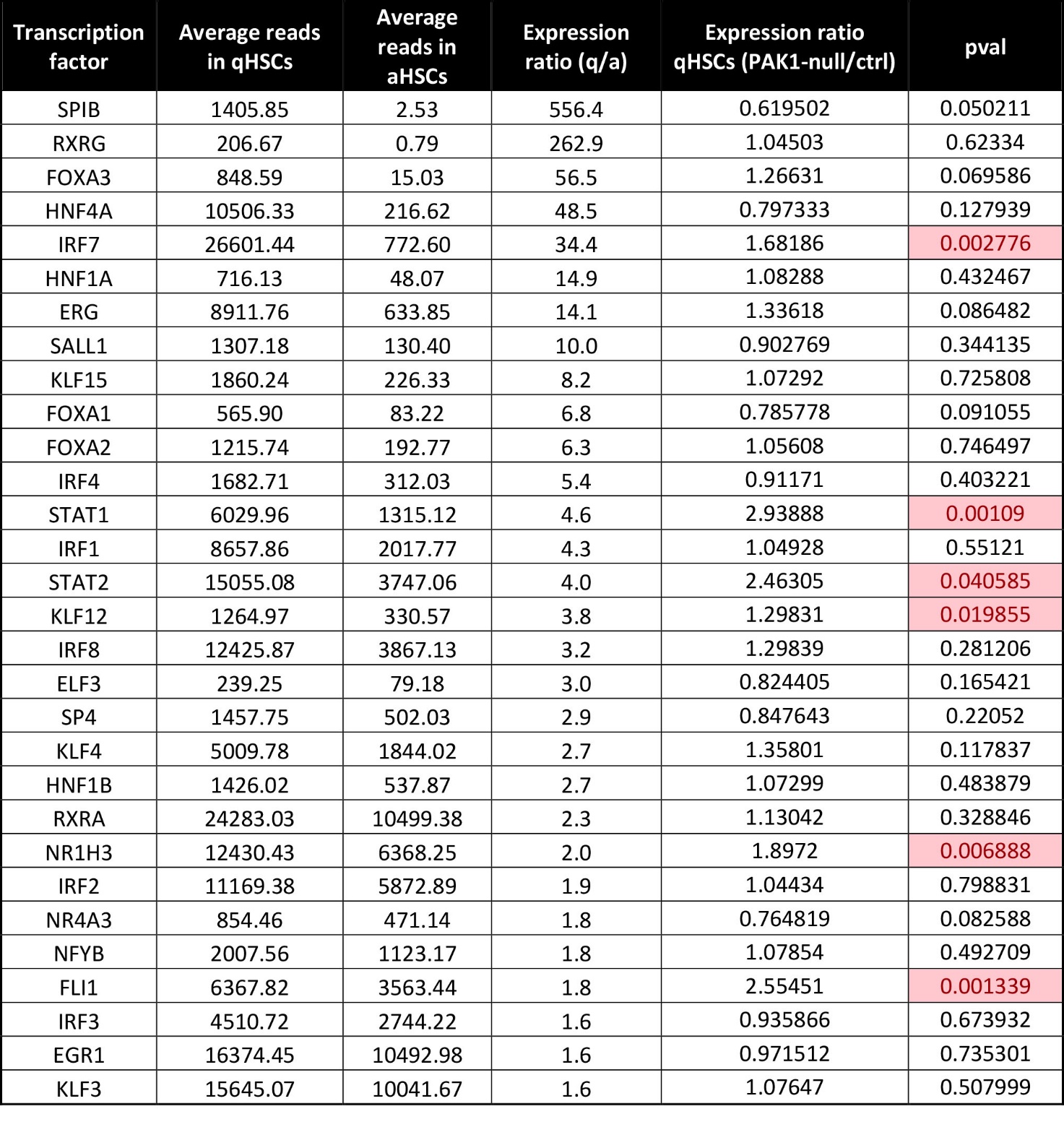

**Supplementary Table 1.** TF motifs enriched in OQUQ promotor peaks; >100 reads RNA expression in q; q/a expression ratio >1.5.

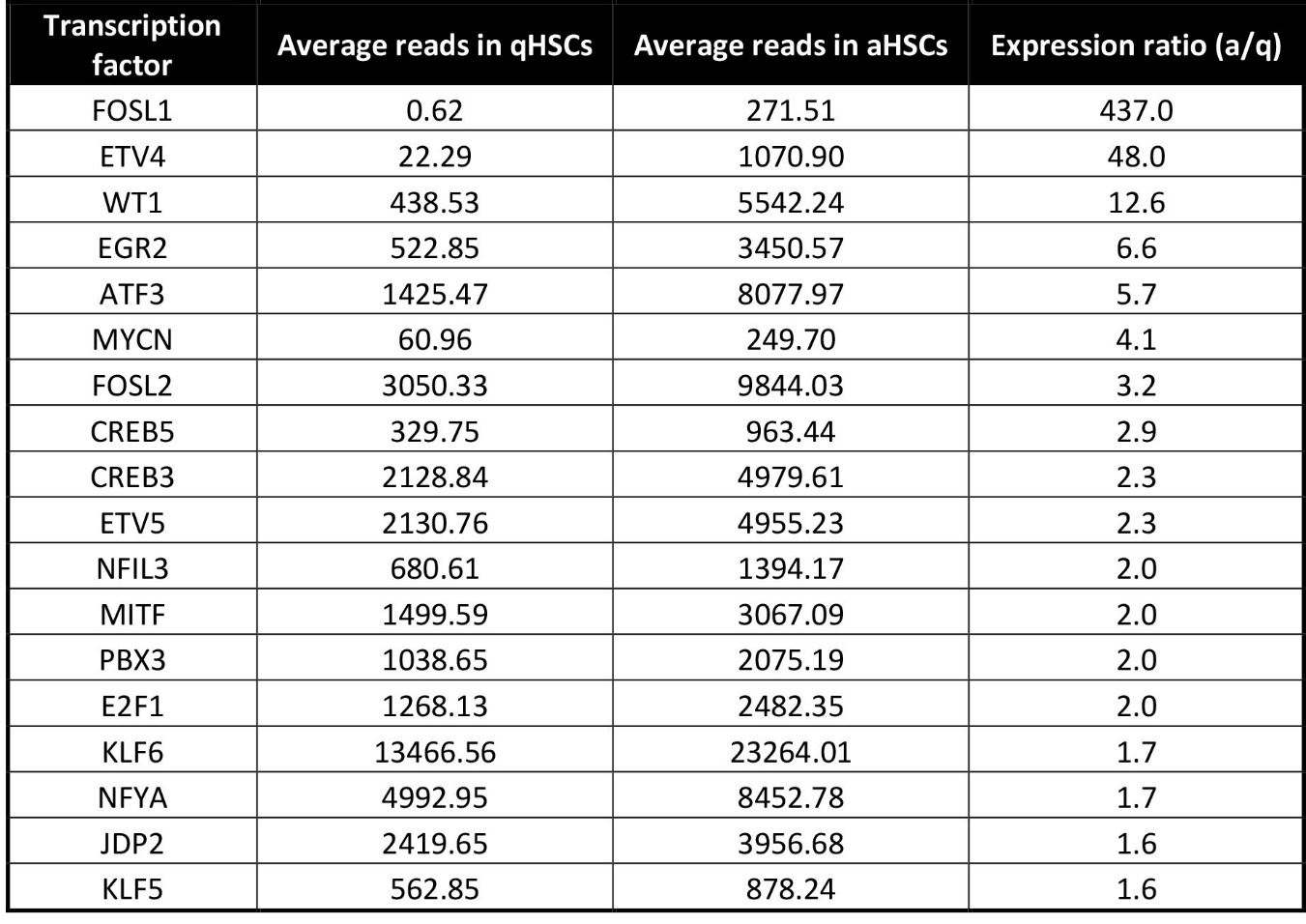

**Supplementary Table 2.** TF motifs enriched in OAUA gene promotor peaks; >100 reads RNA expression in a; a/q expression ratio >1.5.
